# A functional trait approach reveals the effects of landscape context on ecosystem services provided by urban birds

**DOI:** 10.1101/2022.02.28.482331

**Authors:** Timothy M. Swartz, Jason M. Gleditsch, Jocelyn E. Behm

## Abstract

Land use intensification in urban areas can have profound effects on biological communities that provide valuable ecosystem services to urban residents. We used a functional response and effect trait approach to determine how bird species’ responses to local and landscape-scale habitat of urban green spaces affects the supply of cultural and regulating ecosystem services. We sampled bird communities and habitat variables in urban green spaces that varied in local and landscape-scale habitat composition and compiled a dataset of species’ response and effect traits related to nesting, foraging, diet, and visual and acoustic aesthetic appeal. Overall, the landscape-scale context of a green space had a stronger influence on species’ abundances than local-scale habitat. Landscape-scale impervious surface surrounding our study sites interacted with response traits related to nesting in human-built structures, clutch size, and consumption of invertebrates to drive bird species’ abundances. Because correlations between response and effect traits can influence the effect traits available to provide ecosystem services at a site, we explored the correlation of these three response traits to a suite of effect traits and found the response traits were correlated with several effect traits related to diet and regulating services but correlated with few of the plumage and acoustic traits that produce cultural services. Finally, we found that effect traits associated with cultural and regulating ecosystem services varied strongly along the landscape-scale gradient of urbanization. Sites with high impervious surface cover are expected to have low levels of invertebrate pest control and visual appeal but high levels of acoustic appeal, diet evenness (generalism), and granivory. Overall, our study highlights the key role of landscape-scale habitat in driving bird-mediated ecosystem services and underscores the importance of regional urban planning to create healthy and livable cities.

## Introduction

By 2030, 60% of the Earth’s human population is expected to live in cities (United Nations 2018) and global urban land area will have tripled from its 2000 baseline (Seto et al. 2011, Seto et al. 2012). In this context, it is increasingly important for the green spaces within cities to provide a wide range of high-quality ecosystem services to urban residents (Young 2010, Aronson et al. 2017, Dickinson and Hobbs 2017). As such, understanding how the species within urban green spaces generate ecosystem services has become a research priority (Ziter 2016, Schwarz et al. 2017).

Theoretically, the contribution of a particular species to an ecosystem service can be predicted by its “effect traits” – attributes of a species that influence its effect on an ecosystem service (Lavorel and Garnier 2002, Suding et al. 2008). When the relationships between effect traits and ecosystem services are well-established, greenspaces can be managed to provide particular ecosystem services by selecting species with optimal effect traits. This functional trait approach has shown promise for urban green space and ecosystem service management. Plants are typically the focus of these research efforts because strategic plantings can directly increase the abundance of species with optimal effect traits (e.g., Tran et al. 2020, Kleyer 2021). However, application of this functional trait approach to other organisms, like animals, which also provide numerous ecosystem services (e.g., Whelan et al. 2008, Ghanem and Voigt 2012, Valencia-Aguilar et al. 2013), poses distinct challenges since they cannot easily be introduced directly into urban green spaces. Rather, ecosystem services provided by animals depend on whether habitat characteristics within and around the green space attract and support species that contribute to the service. As a result, “response traits” that affect a species’ tolerance of environmental conditions (Lavorel and Garnier 2002, Suding et al. 2008) must also be considered.

Species with response traits suited to habitat conditions are expected to be more abundant in a particular locality (e.g., Cane et al. 2006, Luck et al. 2013, Pardo et al. 2018). Response traits may interact with both local-scale habitat within a green space as well as a green space’s landscape-scale context (i.e., the habitat surrounding it). Local scale habitat suitability can depend on attributes like vegetation structure and composition that relate to how green spaces are managed (Aronson et al. 2017). The extent of suitable habitat in the landscape surrounding a green space can also have a marked effect on species abundances within a greenspace (e.g., Blair 1996, Litteral and Shochat 2017). From a management perspective, it is particularly important to determine the relative influence of local-versus landscape-scale habitat variables on the species in a green space. If species respond more strongly to the landscape-scale variables that are beyond a green space manager’s jurisdiction, coordinated urban planning at a regional scale may be necessary to augment biodiversity and the services it provides.

Consequently, the provisioning of ecosystem services in urban green spaces by animal species is a result of correlations between species’ response and effect traits. Species with the optimal effect traits that provide desired ecosystem services will be abundant in a green space only if they have response traits that allow them to tolerate the site’s habitat conditions. Relationships between response and effect traits will drive variation in effect trait distributions across an urban landscape along gradients in the habitat variables to which species are responding. As a result, due to their habitat conditions, green spaces may vary significantly in the amount of ecosystem services they provide (Gardiner et al. 2013, Aronson et al. 2017). While relationships between response and effect traits are predicted (Suding et al. 2008, Stachewicz et al. 2021), explorations of these relationships and their influence on effect trait variation in the context of ecosystem service provisioning in urban landscapes are scant. Further, it is unknown how these response-effect trait relationships vary for different types of ecosystem service.

Two important groups of ecosystem service in urban areas that involve animals are regulating and cultural ecosystem services (Millennium Ecosystem Assessment 2005). Regulating services, like seed dispersal and biological control of pests, tend to depend on effect traits related to trophic position, foraging strategy, and morphology (Luck et al. 2012). Although less well understood, animals also provide numerous cultural ecosystem services, which encompass the diverse range of non-material benefits people derive from nature, like spiritual, psychological, or recreation benefits (Millennium Ecosystem Assessment 2005). Effect traits underlying cultural services include those that influence how humans perceive or interact with species (Goodness et al. 2016, Echeverri et al. 2019a), such as traits relating to behavior and aesthetic appeal. Cultural services can be particularly important in cities due to limited opportunities for human-nature connections (Goodness et al. 2016, Dickinson and Hobbs 2017).

Here we use response and effect functional traits to examine how habitat conditions impact the regulating and cultural ecosystem services provided by bird communities in urban green spaces. Birds readily occupy habitats throughout the urban landscape (Blair 1996, Callaghan et al. 2020) and are sufficiently well-studied that detailed trait data can be obtained across a broad suite of response and effect traits (e.g., Wilman et al. 2014), from diet and foraging behavior to plumage coloration and song characteristics. In turn, these traits link them to a wide range of regulating and cultural ecosystem services (Sekercioglu 2006, Whelan et al. 2008, Echeverri et al. 2019a, Cameron et al. 2020). We identify the important response traits that are driving variation in bird species abundances across local and landscape habitat variables. We then quantify response-effect trait correlations and explore how effect trait composition varies with local and landscape habitat conditions. We conclude by modeling how the supply of cultural and regulating ecosystem services may vary across urban green spaces due to these habitat and trait relationships. Together, these analyses create an integrative approach exploring how urban landscapes and green spaces can be managed for bird-mediated ecosystem services.

## Methods

### Study Region

We studied public urban green spaces in Philadelphia County, Pennsylvania, USA. As the sixth most populous city in the United States (U.S. Census Bureau 2019), Philadelphia’s green spaces service a population of 1.58 million people. These green spaces consist of about 4,100 hectares of public parks including plazas, playgrounds, sports fields, and forest preserves (Philadelphia Parks & Recreation 2021) as well as about 1,000 hectares of approximately 40,000 vacant lots (Pearsall 2017, City of Philadelphia 2020). The Pennsylvania Horticultural Society has renovated 12,000 of these vacant lots (South et al. 2018), which now function as supplemental public green space in many neighborhoods (Heckert and Kondo 2018).

### Study Site Selection

We measured bird communities and habitat at 60 sites in green spaces across the urban landscape of Philadelphia (Table S1). We selected study sites to be distributed across two gradients representing the availability of suitable habitat at the local and landscape scales (after Cox et al. 2018): 1) *local scale tree canopy cover* within the site boundaries and 2) *landscape scale cover of impervious surfaces,* which includes roads, sidewalks, and buildings within a 500m radius from the site centroid (Weaving et al. 2016). Both gradients were derived from a high-resolution land cover raster (3m resolution; City of Philadelphia 2011) using ArcMap 10.7.1 (ESRI, Redlands, California, USA). Selected study sites were located ≥250 m from each other to limit double-counting individual birds (after Ralph et al. 1995). The selected study sites were located within green spaces ranging from small vacant lots (0.05 ha) to large forest preserves (e.g., Wissahickon valley park, 826 ha; Table S2).

### Avian Point Counts

We sampled bird communities using 5-minute, 50m radius point counts at each site. All birds seen or heard were counted and identified and their distance from the observer was recorded with a laser range finder (Impact 850, Vortex Optics, Barneveld, WI). Counts were conducted at the center of small sites (<100m across) or at a random point ≥50m from the edge of larger sites. Sites were sampled during the breeding season of resident birds (May 21 – July 29, 2019). To reduce bias caused by factors that influence the detectability of birds (Buckland et al. 2012), all counts at all sites were conducted by a single observer (TMS) from 6:00 AM to 10:30 AM, when bird activity and detectability are highest (Rega-Brodsky and Nilon 2017) and no counts took place during periods of precipitation or high wind. We sampled each site at two-week intervals and varied the daily sampling order of sites. Each site was visited at least four times, with 55 of the 60 sites being visited 5 times. We calculated site-level species abundances as the mean number of individuals per species seen or heard across all sampling visits rounded up to the nearest integer to accommodate use of ‘fourth corner’ analyses (see below).

### Landscape Context and Local Habitat Variables

We measured landscape- and local-scale variables for each site to determine how they were related to bird response traits. At the landscape scale, we measured landscape context based on proportion of impervious surface cover (3m resolution; City of Philadelphia 2011) and tree canopy cover (3m resolution raster generated from the 2018 Philadelphia Tree Canopy Assessment; see O’Neil-Dunne 2019) within a 500m buffer radius of the avian sampling point in ArcMap (Table 1). Other measures of landscape composition and configuration surrounding the sites calculated with the R package *landscapemetrics* (Hesselbarth et al. 2019) were considered for inclusion in the analysis but were highly correlated with the proportion of impervious surface cover and were therefore excluded.

**Table 1.**
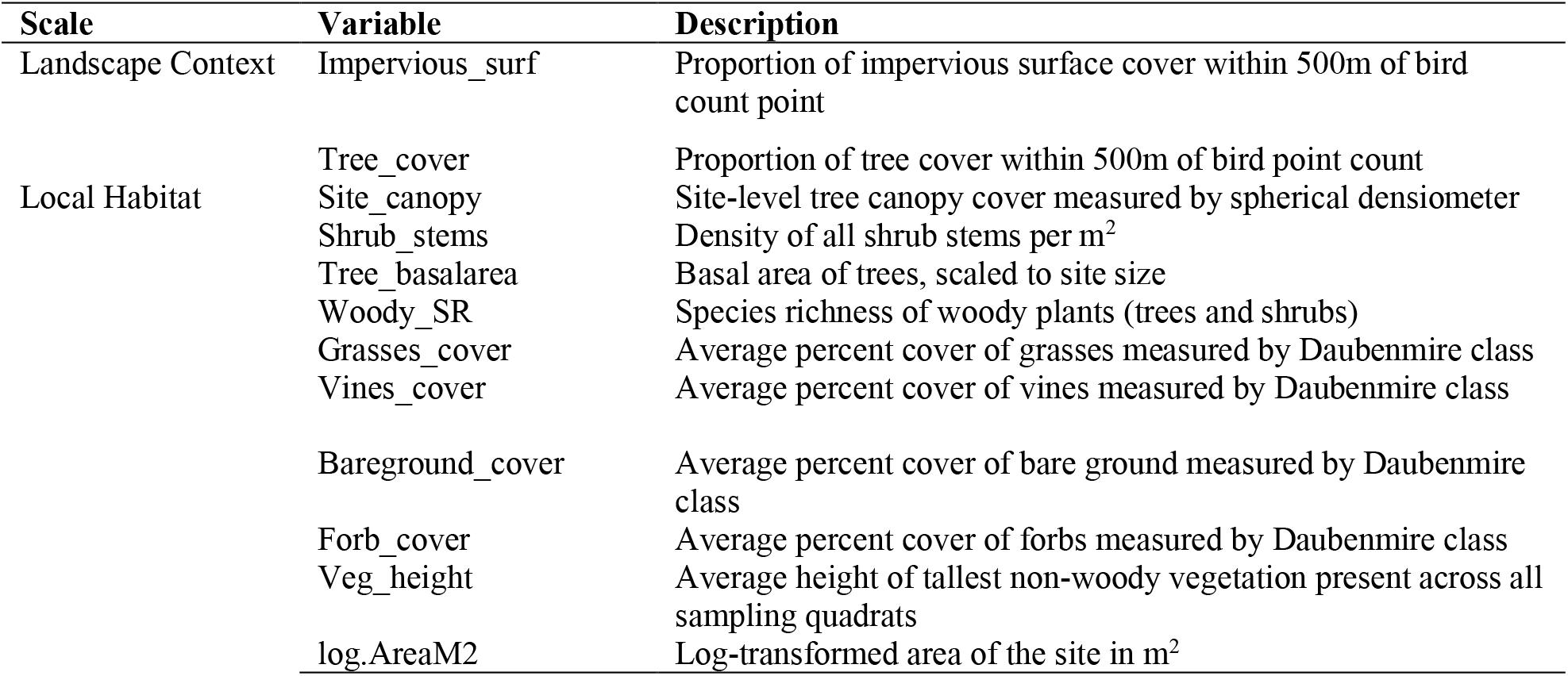
Habitat variables assessed for each study site.

At the local scale, we assessed local habitat using field surveys of herbaceous and woody vegetation composition and structure (Table 1) at each site. We modified the Breeding Biology Research and Monitoring Database protocol (BBIRD; Martin et al. 1997) to measure woody vegetation and used Daubenmire (Daubenmire 1959) methods to measure herbaceous vegetation. There was substantial variation across sites in terms of the density and distribution of woody vegetation (trees and shrubs), so we used two sampling methods. A “complete census” method was used for sites (N = 33) with sparse enough vegetation that all trees and shrubs could be censused within a 100m radius from the bird point count location or within the site boundaries for sites smaller than 100m in radius. For sites with higher woody plant density (N = 27) we used a “census plot” method based on the BBIRD field protocol, which is designed for assessing relatively homogenous habitats (Martin et al. 1997). For this method, woody vegetation was assessed in four circular plots, with one plot centered on the point count location and three located 10–30 m (depending on the size of the site) from the center point and evenly arranged at 120° angles. The radius of the census plot was adjusted based on vegetation density to ensure sampling efficiency (reduced from 10m radius to 5m radius in very dense shrubs; after Martin et al. 1997). For both the complete census and census plot method, we recorded the identity of all woody species, counted shrub stems, and measured the diameter at breast height (DBH) of all trees >1 cm DBH within the whole site (complete census) or circular plot (census plot). We used counts of stems and tree DBH to calculate stem densities and basal area to use in subsequent analyses.

Herbaceous vegetation (forbs and grasses) was sampled using a set of eight paired 0.5 m^2^ quadrats. A pair of quadrats was located within each of the four BBIRD plots in census plot sites or their approximate location for complete census sites. In each quadrat, we visually estimated the percent cover of grass (Daubenmire 1959) and recorded the height of the tallest stem of herbaceous vegetation. The eight quadrat values were averaged for a site-level measurement for each variable. Within each plot we also estimated tree canopy cover using a spherical densiometer (Lemmon 1956) and averaged these four measurements for a site-level canopy cover value.

### Response and Effect Traits Acquisition

For each species observed during field sampling, we compiled a dataset of 16 response traits related to how birds respond to environmental conditions and 14 effect traits related to regulating and cultural ecosystem services (Table 2). We used response traits related to morphology (body size and shape), diet composition, foraging stratum, and reproduction (nest location and clutch size) (after Luck et al. 2012, Leveau 2013, Callaghan et al. 2019). For effect traits contributing to cultural services, we used acoustic traits related to song complexity or variability, as well as bird size, shape, and plumage color. For effect traits contributing to regulating services, we used diet traits.

**Table 2.**
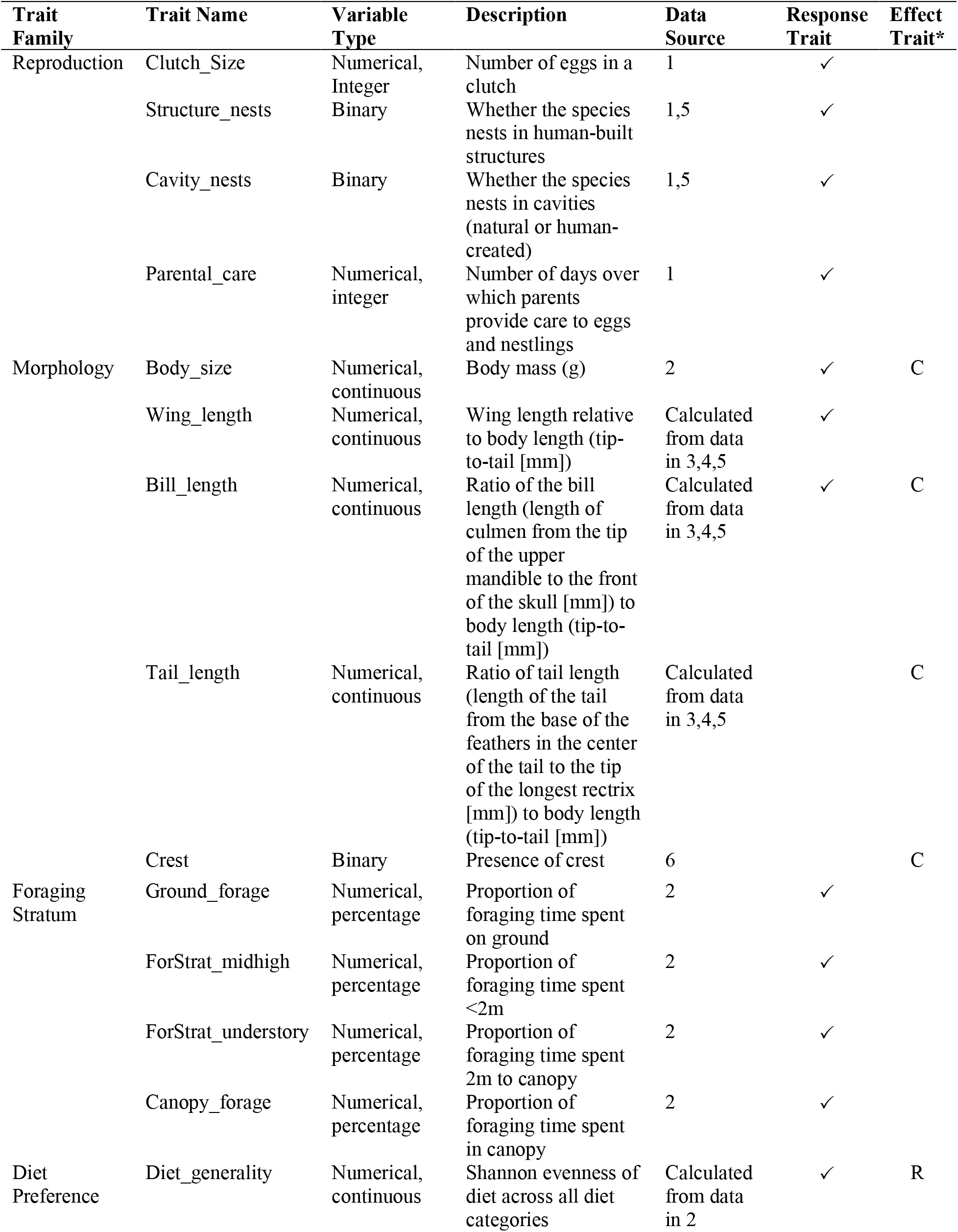
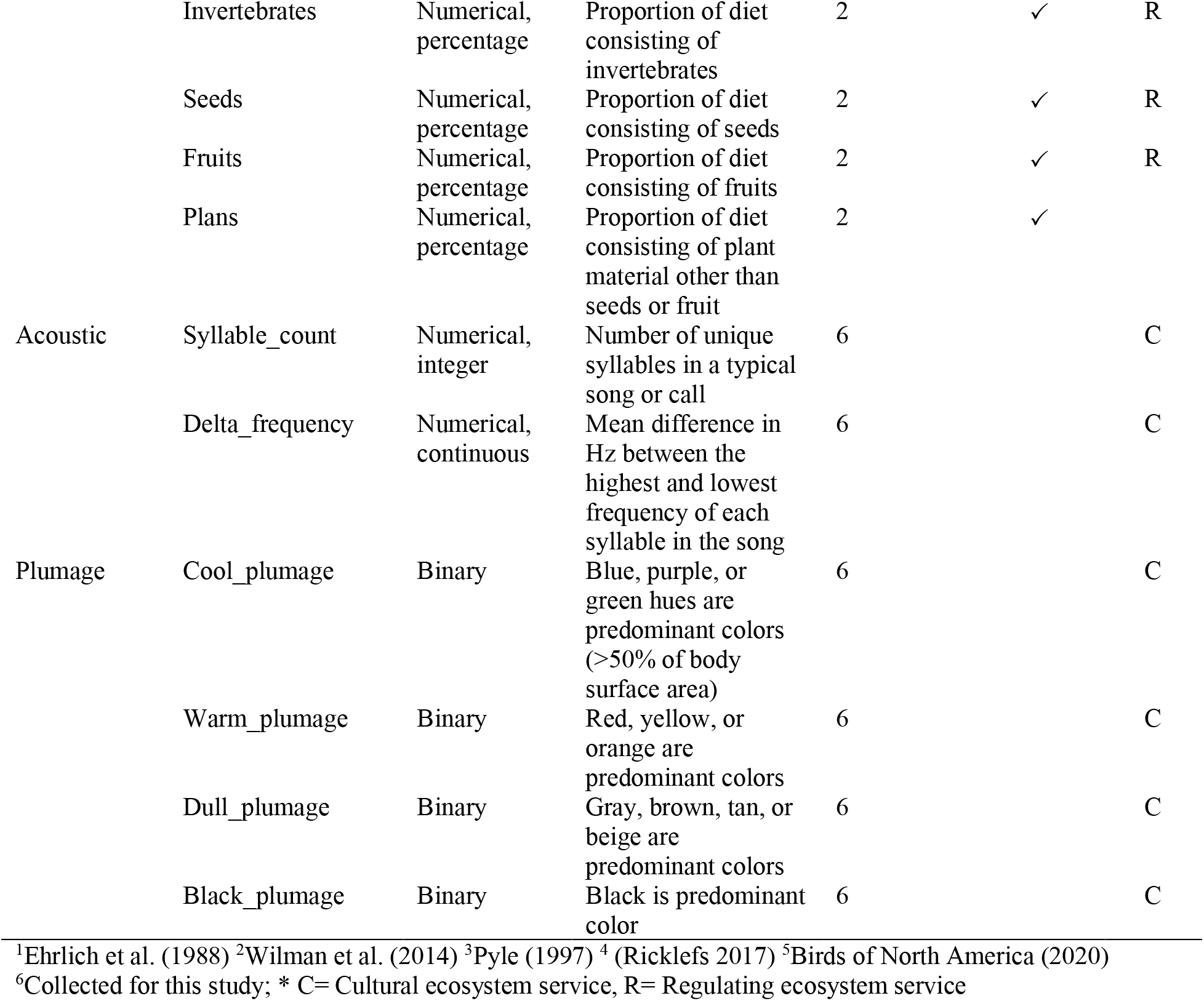
Descriptions and data sources for bird response and effect functional traits used to explain how local habitat and landscape context shape bird communities and ecosystem services in urban green spaces.

For all response and effect traits apart from acoustic traits, we obtained trait values from the literature (sources detailed in Table 2). Due to the absence of published acoustic trait data, we measured acoustic traits related to song variability and complexity in Raven Pro (v. 1.6.1, The Cornell Lab of Ornithology, Ithaca, NY). Following Echeverri et al. (2019a), we selected a single representative recording for each species from the Xeno-Canto database (www.xeno-canto.org). We selected recordings with low background noise, consisting of the primary song, and recorded from southeastern Pennsylvania or the surrounding region, where possible (Echeverri et al. 2019a). A call was used for species lacking song-type vocalizations (N = 12). The full list of recordings used is available in Table S3. For species with very large song repertoires (e.g., Northern Mockingbird [*Mimus polyglottus*]), measurements were confined to a representative 30 s clip. Two song characteristics were measured: number of syllables and total song frequency range (Table 2). Research explicitly linking acoustic traits to human perception of bird song aesthetics is scarce (Goodness et al. 2016), so we measured traits that capture the variability and diversity of the pitch and syllables, which may reflect how acoustically interesting and pleasant the songs are to human listeners (Echeverri et al. 2019a).

### Statistical Analyses

We performed a series of analyses to link habitat and ecosystem services with response and effect traits of bird species in urban green spaces. We provide an overview of our analytical approach here with a detailed description of our methods to follow. We first identified response traits and local and landscape habitat variables driving the abundances of bird species across our study sites. We then determined whether any of these response traits were correlated with effect traits that drive ecosystem service supply, suggesting a link between habitat variables and ecosystem services. To explicitly examine the links between effect traits, habitat, and ecosystem service supply at the community scale, we assessed how the summed effect traits in bird communities varied with habitat variables. Then we assessed how ecosystem services calculated from community-scale effect trait totals were influenced with habitat variables. All analyses were conducted in R (R version 4.0.0; R Core Team 2015); code and data to replicate our procedures are provided in Code S1 and Data S1.

#### Identifying key response traits and habitat variables

We identified the key response traits driving bird species’ responses to habitat variables using the “traitglm” function in the R package *mvabund* (Wang et al. 2012). This approach models the abundance of species across multiple sites as a function of the species’ traits, environmental habitat variables, and the interaction between traits and habitat (Brown et al. 2014, Warton et al. 2015). The relative magnitude of the coefficients of the trait-habitat interactions are an indicator of their importance in determining bird species abundances at sites (Bartomeus et al. 2018) and are a way to identify the key response traits with respect to habitat variables. Three matrices were used as inputs for this ‘fourth-corner model’ (see Legendre et al. 1997): 1) a site-by-species matrix of bird abundances at each of the 60 sites, 2) a species-by-trait matrix consisting of the trait values for each species, and 3) a site-by-habitat matrix for measurements of local- and landscape-scale variables. The landscape-scale variables were percent cover of impervious surface and tree cover within 500m of our study sites. Local-scale variables were the first two axes derived from a principal components analysis (PCA) performed using the “rda” function from the vegan package in R (Oksanen et al. 2019) on our full set of local habitat variables (see Table 1) to reduce the dimensionality of the data. These two axes captured 52.02% of the variation in our local-scale variables and represent gradients of vegetation density. Specifically, the first axis (PC1 – 34.82% of variation) represents a gradient in woody vegetation density from open lawn sites to natural, complex forest vegetation with high shrub and large tree density. The second axis (PC2 – 17.2%) represents a gradient of herbaceous vegetation density, from those with extensive bare ground to sites with unmown, overgrown herbaceous vegetation (Fig. S1 & S2).

We used the default negative binomial family and specified the “glm1path” method to employ a LASSO penalty to perform model selection and reduce to zero the coefficients of trait-habitat interactions that do not improve model fit as measured by the Bayesian Information Criterion (Warton et al. 2015). We identified important response traits as those with standardized interaction coefficients ranking in the 95^th^ percentile for absolute effect size and used these important response traits in subsequent analyses.

#### Correlations between response and effect traits

Effect traits have no presumed relationship with habitat conditions (Suding et al. 2008). Rather, habitat conditions impact ecosystem services via correlations between species’ response and effect traits. We investigated these relationships by testing for correlations between the important response traits we identified and our 14 effect traits across all species observed in this study using an abundance-weighted linear Pearson correlation (after Pakeman 2011). The mean abundance of each species across all study sites was used for weighting. This allowed us to account for the contribution of each species to the analysis in proportion to their abundance across the whole data set. This procedure was performed with “weightedCorr” function from the *wCorr* package (Emad and Bailey 2017).

#### Community-scale effect trait-habitat associations

The ecosystem services produced in a site depend on the entire community of service providers (Suding et al. 2008). Accordingly, we used a redundancy analysis (RDA) to identify associations between habitat variables and community-summed effect traits to determine how effect traits vary across the urban landscape. RDA is a constrained ordination method that explains variation in community data using habitat variables (Ter Braak 1986, Borcard et al. 2018). Here, our community data consisted of a community-summed effect trait matrix where effect trait values were multiplied by each species’ site-level mean abundance then summed for all species (see Table 2 for traits). These community-summed effect traits represent the total amount of effect trait produced by the bird community at each site. The habitat data were a matrix of the two local habitat PC axes (see *Identifying key response traits and habitat variables* above) and the two landscape context variables (see Table 1 for variables). Community-summed effect trait and habitat variables were mean-centered and scaled by the standard deviation. The RDA was performed using the “rda” function in *vegan* (Oksanen et al. 2019). We used the “anova.cca” function, also in *vegan*, to perform permutation tests of the significance of the overall ordination and the significance of each RDA axis. An adjusted R-squared value was generated with the “RsquareAdj” function as a measure of goodness of fit for the overall ordination solution.

#### Modeling ecosystem service supply

To quantify how the habitat-effect trait associations scale to affect ecosystem services, we used hypothesized trait-ecosystem service relationships to calculate seven ecosystem service scores from the community-summed effect traits (after Tran et al. 2020; Table S4). In short, we calculated simple additive measures for each effect trait family (visual, acoustic, and diet) by summing the scaled values for each effect trait at each site, with each trait either contributing positively or negatively to a service, based on the literature (Table S4). We generated linear models to assess the relationship between the supply of each ecosystem service quantified from the model and landscape scale impervious surface cover. We adjusted our alpha level to account for multiple tests (N = 7) using a Bonferroni correction (alpha = 0.007).

## Results

Across our 60 sites, 3,607 individuals from 45 bird species were observed and included in our analyses. Mean site-level species richness was 8 (SD = 3.5, range = 2 – 21). The three most common species, which comprised 66.7% of all individuals observed, were House Sparrow (*Passer domesticus*; 38.5%), European Starling (*Sturnus vulgaris*; 17.2%), and American Robin (*Turdus migratorius*; 11.0%). The average site-level abundance of individuals per visit was 8.6 (SD = 5.4, range = 1 – 52).

### Identifying key response traits and habitat variables

Overall, trait-habitat interactions were stronger for landscape than for local habitat variables; the average magnitude of interaction coefficients for response traits at the landscape scale was 0.092 (range −0.172–0.176), which was nearly three times that of the interaction coefficients for the local scale (0.034, range −0.05–0.085; Fig. 1). We identified three important response traits with trait-habitat interactions in the 95^th^ percentile across all variables and each of these response traits was associated with landscape-scale impervious surface cover. Species that nest in human-built structures, species with smaller clutch sizes, and species with lower proportion of invertebrates in their diets were more abundant in sites with higher landscape-scale impervious surface cover (Fig. 1).

**Figure 1.**
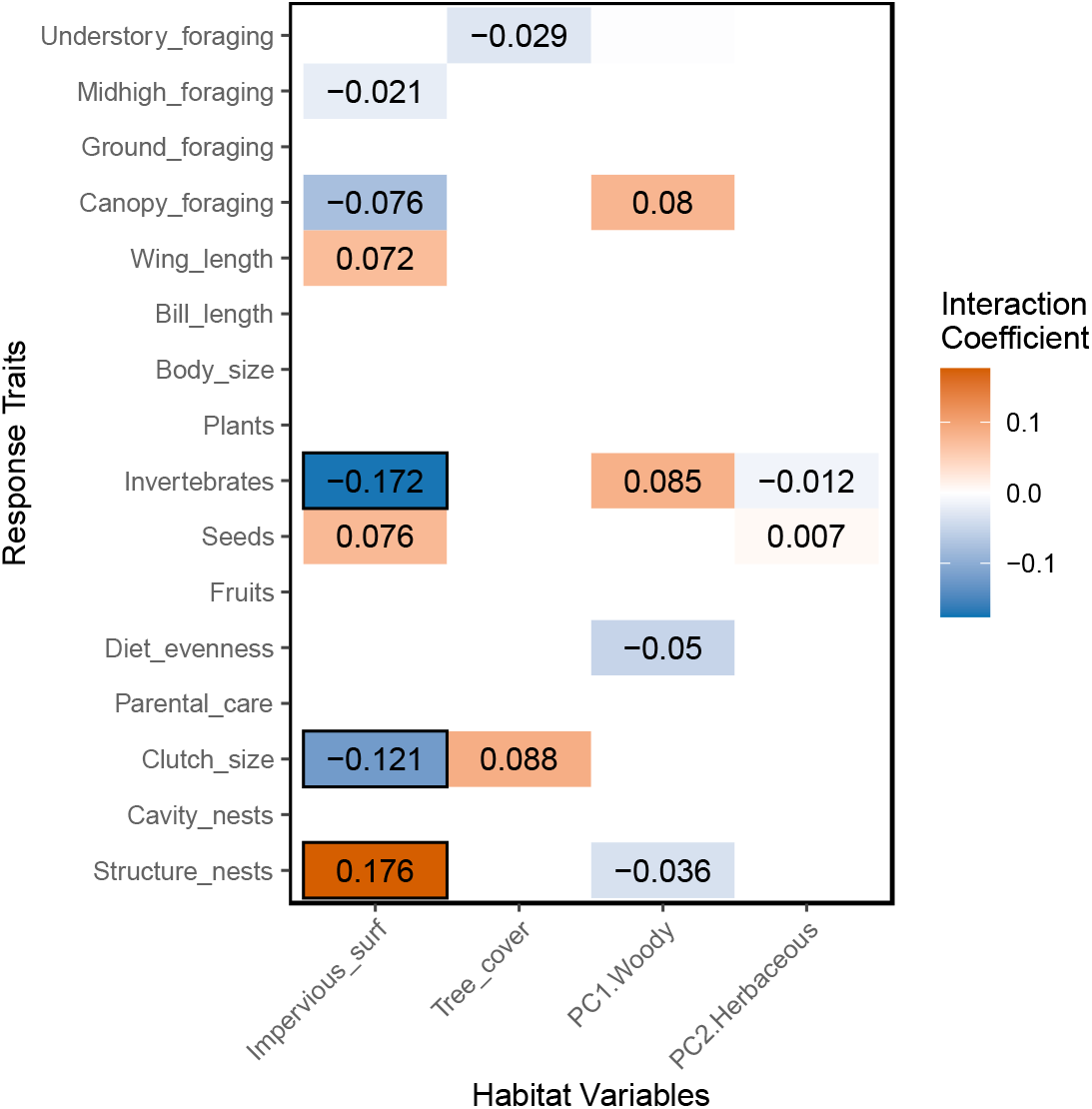
Tile plot indicating the direction and strength of interaction coefficients between avian response traits and landscape context (Impervious_surf and Tree_cover) and local habitat (PC1.Woody and PC2.Herbaceous) variables from multiple linear regression modeling. Interactions without sufficient statistical support appear as blank tiles. Black boxes highlight the strongest response trait-environmental variable interactions (magnitude of coefficients in the 95^th^ percentile across all variables).

### Correlations between response and effect traits

The abundance-weighted Pearson correlation showed that the three important response traits we identified (i.e., invertebrate diet, clutch size, and nesting in built structures) were each significantly correlated to several effect traits related to regulating and cultural ecosystem services. (Fig. 2). Nesting in structures and invertebrate diet correlated with each of the diet effect traits, except diet evenness. Structure nesters were more likely to consume seeds rather than invertebrates or fruits. Invertebrate consumers tended to also be fruit eaters and not seed eaters. For cultural ecosystem service traits, response traits were only correlated to shape traits. Species with larger clutches tended to be those with lesser body mass and longer bills and those that nest in structures tend to have shorter tails.

**Figure 2.**
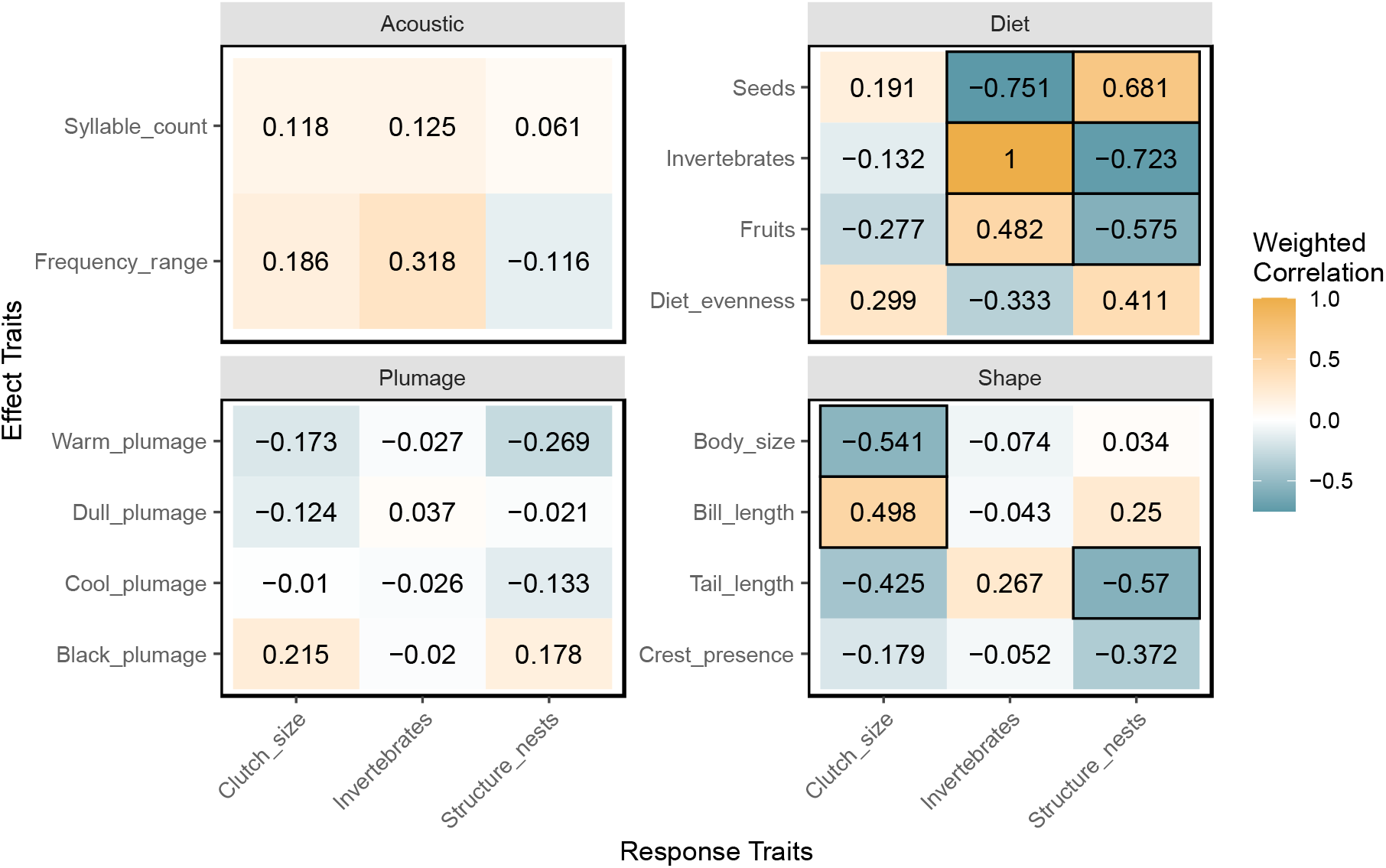
Tile plots of pairwise abundance-weighted Pearson correlations between avian response and effect traits, grouped by effect trait family. Acoustic, plumage, and shape families contribute to cultural ecosystem services whereas diet traits contribute to regulating services. Significant correlations (alpha = 0.05) are outlined with a black border.

### Community-scale effect trait-habitat associations

Our ordination of site-level community-summed effect trait values revealed several links between effect traits and habitat variables (Fig. 3). Overall, the explanatory local and landscape habitat variables significantly captured variation in the effect traits (F_4,55_ = 5.49, *p* < 0.001; adjR^2^ = 0.23) and multicollinearity among explanatory habitat variables was low (VIF < 5 for all). Together, the habitat variables explained 28.55% of the total variance in community-summed effect traits in our redundancy analysis (RDA). Permutation tests indicated that only the first RDA axis explained a significant amount of variation in community-summed trait values (RDA1: F_1,55_ = 19.07, *p* < 0.001; 24.78% of total variance explained) and the only explanatory variable that had a significant relationship with RDA1 was impervious surface (i.e., Impervious_surf; RDA1: F_1,55_ = 18.12, *p* < 0.001), which had a negative relationship with RDA1. RDA2 explained substantially less variation in summed trait values and was not significant (F_1,55_ = 1.44, *p* = 0.67; 1.87%).

**Figure 3.**
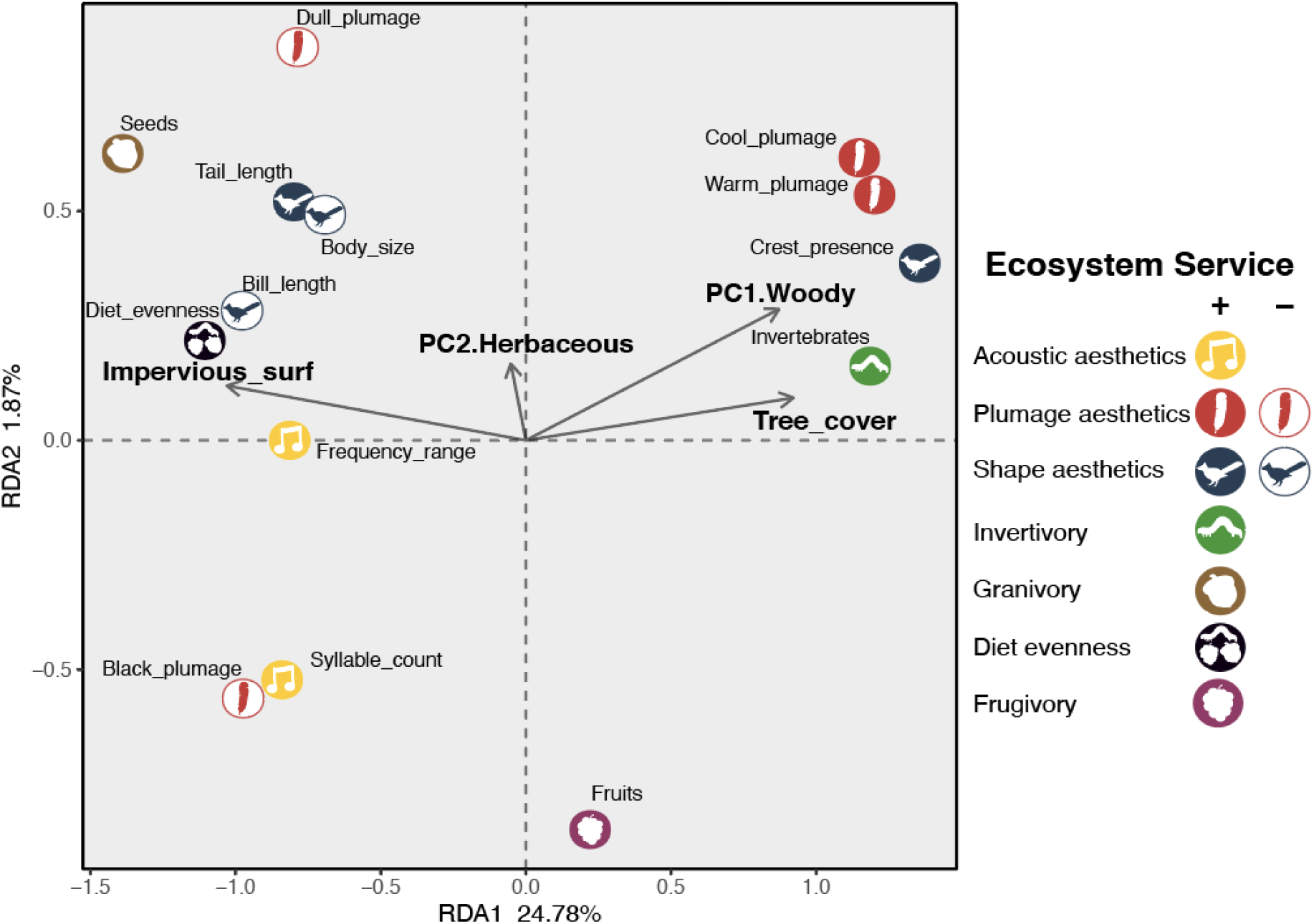
Redundancy analysis (RDA) biplot showing the centroids of avian effect traits as explained by environmental variables (vector arrows, bold text). Effect traits are grouped by the ecosystem service to which they contribute (symbol and color) as well as the direction of their contribution to the service (positive or negative, see Table S3). Note, RDA axes displayed on different scales.

For the most part, effect traits that contribute positively to the supply of plumage- and shape-related aesthetic services (filled red and blue symbols in Fig. 3) were more prevalent at sites with lower landscape impervious surface cover (positive relationship with RDA1) (Fig. 3). In contrast, traits that supply acoustic services (filled yellow symbols) were more prevalent at sites with high impervious surface cover (negative relationship with RDA1). For regulating services, relationships between community-summed effect traits and RDA1 varied by service, with a negative relationship with RDA1 for granivory and diet evenness and a positive one for invertivory.

### Modeling ecosystem service supply

These relationships suggested by the RDA were also observed in our ecosystem service scores (Fig. 4). Services associated with diet evenness, granivory, and acoustic aesthetics are predicted to increase with impervious surface cover, whereas services related to invertivory, plumage, and shape aesthetics are expected to decrease. Frugivory services did not strongly correspond to impervious surface levels.

**Figure 4.**
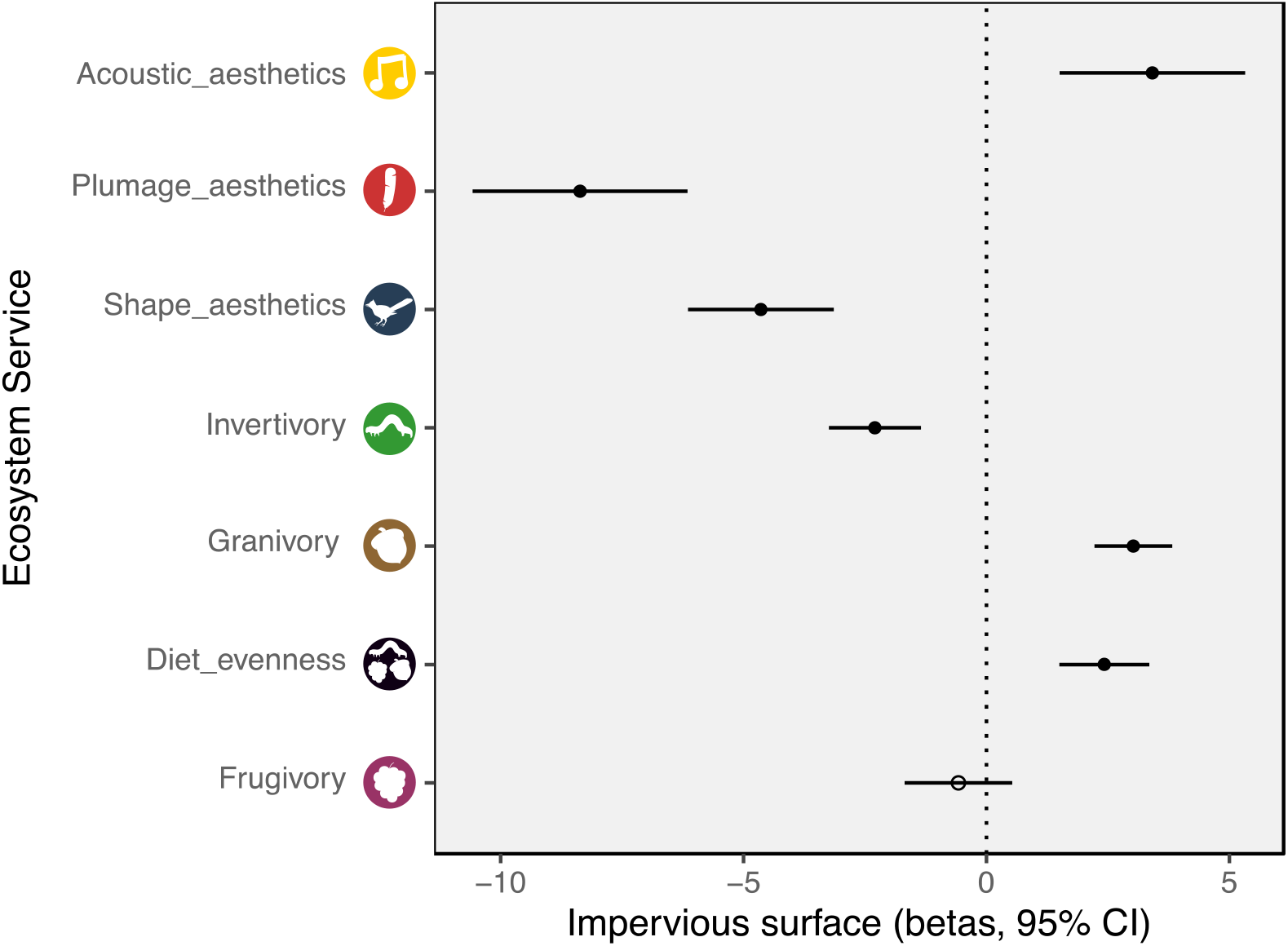
Plots depicting the effect of four local and landscape habitat variables on 7 ecosystem service scores. Standardized model coefficients and 95% confidence intervals from linear regression models are displayed, filled circles denote statistically significant coefficient (alpha = 0.007).

## Discussion

In this study, we applied a functional trait approach to understand how the response of bird species to local and landscape habitat characteristics shape the community-scale supply of ecosystem services provided in urban green spaces. Most urban animals and their associated services cannot easily be managed through direct introductions of species into green spaces. Instead, we show how response and effect functional traits can be used to identify habitat characteristics important for supporting urban bird communities that provide valued ecosystem services for human well-being. Although all of our sites are located within the boundary of Philadelphia’s city limits, bird species still responded strongly to the habitat gradient generated by land use intensification, which has implications for the provisioning of ecosystem services. In our study, landscape-scale impervious surface cover influenced a range of functional traits and thus affects the availability of both regulating and cultural ecosystem services provided by bird communities. In light of the ongoing expansion and intensification of urban land cover (Seto et al. 2011, Seto et al. 2012), these findings have important ramifications for urban ecosystem service management.

A major component of managing urban green spaces for ecosystem services entails identifying interactions between habitat and species’ response traits, since these interactions affect the presence and abundance of potential service-providing species. Although the urban green spaces we studied varied considerably in their local-scale habitat structure and composition, the strongest trait-habitat interactions were observed for landscape-scale variables, especially impervious surface cover. Thus, the scale at which green space management decisions are made (local) may not match the scale at which bird communities respond to habitat (landscape). The influence of landscape-context on urban bird diversity has been clear for some time (Donnelly and Marzluff 2004, Luther et al. 2008), but local-scale habitat interventions remain a key approach to green space management (Aronson et al. 2017). While beneficial to urban bird conservation (e.g., Threlfall et al. 2017, Reynolds et al. 2019), these local-scale efforts may have limited impact on ecosystem services that are affected by species’ abundances. Some green spaces surrounded by extensive impervious surface cover may simply be too small or isolated to support sufficiently large populations of service providers. Local-scale efforts to protect species of conservation concern should be coupled with landscape-scale planning to address ecosystem service needs (Jokimäki et al. 2018, Liordos et al. 2021).

Strong correlations between response and effect traits are considered to render ecosystem services vulnerable to environmental change (Suding et al. 2008, Stachewicz et al. 2021). However, these correlations are crucial to managing habitat for ecosystem services. If effect traits do not correlate with the response traits that interact with habitat to drive species abundances, then effective, predictable habitat management will be elusive. Our abundance weighted response-effect trait correlation analysis revealed connections between response traits and some, but not all, effect traits. For example, structure-nesting correlated with diet effect traits, highlighting the direct connection between the proliferation of anthropogenic nest sites in areas with high impervious surface cover and the supply of diet-based regulating services. In contrast, only weak correlations were observed between response traits and plumage and acoustic effect traits, leaving us without a firm explanation for the observed effect of impervious surface on cultural services. It is possible that correlations between cultural effect traits and response traits do not exist (Suding et al. 2008). However, it seems more likely that either correlated response traits were not selected in the trait-habitat analysis, which identified a limited subset of traits as important, or that correlated effect traits were not included in our analyses. Investigation of effect traits related to cultural ecosystem services is a relatively new endeavor (e.g., Echeverri et al. 2019a, Zoeller et al. 2020) that may reveal as yet unidentified effect traits for which there are correlations. Until such relationships are known, predictable management of cultural services through local and landscape-scale interventions may be difficult.

Our redundancy analysis and ecosystem service models showed how trait-habitat associations scale up to influence ecosystem services at the community scale. Landscape cover of impervious surface was clearly associated with community-summed effect trait values for the diet, plumage, and shape trait families. Notably, we found that the services that granivores could provide, like the consumption of carbohydrate-rich littered food waste (Youngsteadt et al. 2015), may be relatively high in sites with high landscape impervious surface cover. In contrast, invertebrate pest control is likely low. In some respects, this pattern may match demand for such services, with a higher need for litter removal in population-dense areas. However, invertebrate pest outbreaks often occur in urban areas and isolated green spaces within a high impervious surface context may be more vulnerable to such outbreaks (Long and Frank 2020). Furthermore, despite particularly high demand for cultural ecosystem services in areas with more impervious surface cover (Goodness et al. 2016, Valente-Neto et al. 2021), they are likely most scarce there due to the absence of colorful species. By increasing natural cover in the landscape context surrounding urban parks, managers may be able to attract more visually appealing species and improve cultural service supply.

Our functional trait approach revealed that acoustic aesthetics may be one cultural ecosystem service that is generally higher in green spaces surrounded by high impervious surface land cover. While some abundant urban birds (including House Sparrow [*Passer domesticus*]) possess vocalizations consisting of simple, repeating syllables, European Starling (*Sturnus vulgaris*), another species abundant in our study sites, has an extensive song repertoire, counterbalancing the simple House Sparrow song in our community-scale analyses. Other research approaches have shown contrasting results. In one of the few studies to explore the influence of House Sparrow songs on human perception of the urban soundscape, not only were their vocalizations found to be the least appealing but they reduced overall soundscape appeal when their songs were added to a multi-species chorus (Hedblom et al. 2014). Given the current paucity of research on the relationship between bird song and ecosystem services (Goodness et al. 2016), this is an area where future study of the links between traits and human perception in real-world contexts is warranted.

The need to understand how animal traits generate ecosystem services at a community-scale applies to aesthetic traits and to cultural services more broadly. The links we have made between visual and acoustic traits and the cultural services supplied by green space bird communities are reasonable given present evidence (Goodness et al. 2016), but some are untested. We represented these relationships with unweighted additive models since existing research has focused on directionality of trait contributions to aesthetic appeal and on rankings of appeal across species (Lišková and Frynta 2013, Lišková et al. 2015, Echeverri et al. 2019b). Describing the relative importance of and interactions between different traits is needed to develop more complex models. Such models must also account for the scale at which humans experience birds (see Cox et al. 2017). For example, bird species with highly appealing traits that are rare or difficult to observe may make an overall smaller contribution to ecosystem services than less appealing species that are more abundant and gregarious, but these relationships likely vary depending on the human observers involved (e.g., birdwatchers vs. non-birdwatchers; Echeverri et al. 2019a). Research that builds on species-level studies of bird aesthetics to link communities to cultural service supply (Hedblom et al. 2014), would greatly advance our understanding of how global trends in urbanization and green space management will influence the cultural services provided by urban birds.

In providing a step toward functional trait approaches to managing urban green spaces, our study highlights several future avenues for investigation. First, we expect services to shift with the temporal dynamics of bird communities (e.g., Leveau 2022) including the loss of migratory breeding species in winter and the influx of transitory non-breeding migrants in spring and fall (Graves et al. 2019). While breeding bird communities provide a constant supply of cultural services to summer-time park visitors, non-breeding migrants may provide a strong pulse of services over a short period. Additionally, birds are just one of many urban animal taxa which provide ecosystem services. Other services and their associated communities, like biological control by predatory arthropods (Burkman and Gardiner 2014, Gardiner et al. 2014) or cultural benefits provided by charismatic butterflies (López-Hoffman et al. 2010), may be a more easily be managed through local-scale actions (i.e., mowing, tree planting, seeding native plants). These opportunities should be investigated within a functional trait framework. Applying a trait-based approach to other taxa and seasons would undoubtedly provide further insight into the feasibility of managing local green space habitats for ecosystem services

### Conclusion

Urban green spaces are a centerpiece in cities’ efforts to create thriving urban landscapes. In this endeavor, cities face a difficult task of balancing the demands placed on urban green spaces to provide ecosystem services including carbon storage, urban heat mitigation, recreation, psychological benefits, wildlife habitat, as well as space for cultural enrichment and social interaction (Madureira and Andresen 2014, Aronson et al. 2017, Tran et al. 2020). Our goal here has been to implement an approach using functional traits to quantify the relationships between habitat and the service-providing organisms it supports. We suggest that if managers are equipped with knowledge about community-scale functional responses to management actions, they could pursue targeted interventions to augment desirable services. In this process, tradeoffs among ecosystem services are likely (Haase et al. 2014, Dennis and James 2017) and the decision-making process needs to account for the relative value of the various services in question (Manning et al. 2018). Our approach offers a transparent way to address this challenge by making relationships between habitat, functional traits, and services explicit. Careful, community-engaged management is needed to ensure that green spaces and their biodiversity continue to provide healthy, livable conditions for the world’s growing population of urban residents.

## ACKNOWLEDGEMENTS

We are grateful to Payton Phillips, Matthew Helmus, Nicholas Huron, Rachel Spigler, Amy Freestone, and Brent Sewall for their feedback on earlier versions of this manuscript. We thank Spencer Williams and Rachel Lewandowski for their tireless assistance in the field and Brendan Pang and Kayla Perre for their contributions to the data cleaning and functional trait collation. This work was funded by Temple University and TMS was supported by a Temple University Graduate Fellowship.

