## Supplementary Code and Data for "A functional trait approach reveals the effects of landscape context on ecosystem services provided by urban birds": CodeS1_Swartz et al_2022_GCB.Rmd

````
---
title: "CodeS1 for Swartz & Behm_2022- Bird functional traits and ecosystem services"
output: html_document
editor_options: 
  chunk_output_type: console
---

#Part 1: PCA of sites to generate PC axes

##Loading packages
```{r loading packages pt1}
library(tidyverse) #preparing data, plotting results
library(vegan) #running PCA
library(caret) #for centering on means and scaling
library(ggpubr) #for nice plot themes
```

##Loading Data
```{r load data}
library(readxl)
#setwd()

all.data <- "~/DataS1_Swartz_Behm_2022_bird functional traits and ecosystem services.xlsx"

excel_sheets(all.data)

pcveg.data2 <- read_excel(all.data, sheet = "PCVegData")
sitesdetails <- read_excel(all.data, sheet = "sitesdetails")
```

##Running PCA
```{r run PCA}
library(vegan)

pcveg.data1 <- pcveg.data2 %>%
  mutate(log.AreaM2 = log10(AreaM2)) %>%
  dplyr::select(SiteID, DetailType, log.AreaM2, Tree_cover, Impervious_surf, Site_canopy, Tree_basalarea, Woody_SR,Shrub_stems, Veg_height, Forb_cover, Bareground_cover, Vines_cover, Leaflitter_cover, 
                Grasses_cover) 

preObj <- preProcess(pcveg.data1, method=c("center", "scale"))
pcveg.data <- predict(preObj, pcveg.data1[,])

veg.rda <- rda((pcveg.data[,-c(1,2,4,5)]))

vegrda.smry <- summary(veg.rda)

veg.species.scores <- as.data.frame(vegrda.smry$species)
veg.species.scores$Variable <- rownames(veg.species.scores)

veg.species.scores #loadings of environmental variables on each PC axis

veg.site.scores <- as.data.frame(vegrda.smry$sites)

veg.site.scores$SiteID <- (pcveg.data2$SiteID)
veg.site.scores$DetailType <- (sitesdetails$DetailType)

veg.site.scores #PC axis scores for each site
```

```{r pca var explained}
screeplot(veg.rda, bstick = TRUE)
pca.var.expl <- veg.rda$CA$eig/veg.rda$tot.chi
pca.var.expl
```

##PC Plot
```{r pca ordination plot}
library(plyr)
library(ggrepel)

#Generate hulls for Site Types
find_hull <- function(veg.site.scores) veg.site.scores[chull(veg.site.scores$PC1, veg.site.scores$PC2), ]
chulls <- plyr::ddply(veg.site.scores, "DetailType", find_hull)
chulls <- veg.site.scores %>%
  group_by(DetailType) %>%
  slice(chull(PC1, PC2))

#Relevel Site Type factor
veg.site.scores$DetailType <- ordered(veg.site.scores$DetailType,c("UngreenedLot", "GreenedLot", "Plaza", "RecPark", "Preserve"))

chulls$DetailType <- ordered(as.factor(chulls$DetailType),c("UngreenedLot", "GreenedLot", "Plaza", "RecPark", "Preserve"))

site.palette <- c("#0072B2", "#D55E00", "#F0E442", "#999999", "#009E73")
colorBlindGrey8   <- c("#999999", "#E69F00", "#56B4E9", "#009E73", "#F0E442", "#0072B2", "#D55E00", "#CC79A7")
scales::show_col(colorBlindGrey8)

PCA.plot <- ggplot(veg.site.scores) +
  geom_polygon(data=chulls, aes(x=PC1, y=-PC2, group=DetailType, fill=DetailType, color = DetailType),alpha = 0.8)+
  scale_fill_manual(values = site.palette)+
  geom_point(data = veg.site.scores, aes(x = PC1, y = -PC2, color = DetailType)) + 
  labs(fill = "Site Type") + scale_color_manual(values = site.palette)+ 
  geom_text_repel(data = veg.species.scores, aes(x=PC1, y=-PC2,label = Variable),colour = "black", alpha = 0.85)+
  geom_segment(data=veg.species.scores, aes(x=0, y=0, xend=PC1, yend=-PC2), arrow=arrow(length=unit(0.2,"cm")), alpha=0.75, color="gray20")+
  guides(color = FALSE)+
  geom_vline(aes(xintercept = 0), linetype = "dashed",color = "black",alpha = 0.5) +
  geom_hline(aes(yintercept = 0), linetype = "dashed",color = "black",alpha = 0.5) +
  xlab(paste0("PC1  ", round(pca.var.expl[1]*100,2), "%"))+ ylab(paste0("PC2  ", round(pca.var.expl[2]*100,2), "%"))+
  theme(panel.border =  element_rect(colour = "black", fill=NA, size=0.5))+
  theme(panel.background = element_rect(fill = "gray93", linetype = NULL), panel.grid.major = element_blank(),
          panel.grid.minor = element_blank()) + theme(legend.position = "top") + lims(x =c(-2.2,2.2),y= c(-2.2,2.2))

PCA.plot
```

###*Saving plot PDF
```{r saving figure S2}
#pdf(file = "~/FigureS1.pdf",   # The #directory you want to save the file in
    width = 7, # The width of the plot in inches
    height = 7) # The height of the plot in inches
PCA.plot
dev.off()
```

##PC Loadings plot
```{r }
veg.site.scores
veg.species.scores

veg.species.scores
library(cowplot)
PC1.plot <- ggplot(veg.species.scores)+
  geom_bar(aes(reorder(Variable,PC1),PC1),stat = "identity")+coord_flip()+
    theme(panel.border =  element_rect(colour = "black", fill=NA, size=0.5))+
  theme(panel.background = element_rect(fill = "gray93", linetype = NULL),panel.grid.major = 
          element_blank(),panel.grid.minor = element_blank())+
  labs(y = "PC1", x = "Local Habitat Variable")

PC2.plot <- ggplot(veg.species.scores)+
  geom_bar(aes(reorder(Variable,-PC2),-PC2),stat = "identity")+coord_flip()+
  theme(panel.border =  element_rect(colour = "black", fill=NA, size=0.5))+
  theme(panel.background = element_rect(fill = "gray93", linetype = NULL),panel.grid.major = element_blank(),panel.grid.minor = element_blank())+
  labs(y = "PC2", x = "")

PC.loadings.plot <- plot_grid(PC1.plot, PC2.plot, labels = c('a)', 'b)'), label_size = 12)

PC.loadings.plot
```
###*Saving plot PDF
```{r saving PC scoress PDF}
#pdf(file = "~/FigureS2.pdf",   # The #directory you want to save the file in
    width = 6.5, # The width of the plot in inches
    height = 4) # The height of the plot in inches
PC.loadings.plot
dev.off()
```

#Part 2: 'Fourth-corner' multiple linear regression analysis
This is a type of multiple linear regression analysis that is used to identify the response variables that are important for explaining variation in species abundances across our green spaces. This procedure automatically performs model selection to identify the best response trait-habitat relationships.

##Loading packages
```{r loading packages pt2}
library(mvabund)
```

##Load data
```{r load 4th corner data}
library(tidyverse)
library(openxlsx)
library(Rcpp)
library(readxl)
#L Site X Species matrix
L.matrix <- read.xlsx(all.data, sheet = "L.matrix")

L <- L.matrix %>%
  dplyr::select(-SiteID) #need to remove SiteID for fourth corner

#R Site X Environment matrix
R.matrix <- read.xlsx(all.data, sheet = "R.matrix")

R <- R.matrix %>%
  dplyr::select(-SiteID)

#Q Species X Trait matrix
Q.matrix <- read.xlsx(all.data, sheet = "Q.matrix")

Q <- Q.matrix %>%
  dplyr::select(-Species)

#Site X Species Abundance Matrix
birdabundances.all <- read_excel(all.data, sheet = "birdabundances.all")

#Species X Trait Matrix
birdtraits.all <- read_excel(all.data, sheet = "birdtraits.all")
```

##Fitting Fourth Corner Models
```{r fitting fourth corner models}
#fit models----
library(mvabund) #GLMs and fourth corner models using a LASSO penalty

ft2=traitglm(L,R,Q,method="glm1path",family = "negative.binomial")
ft2$fourth #get the values for the fourth corner (traits-environmental relationships)
```

```{r check assumptions}
plot(ft2) #No obvious overdispersion here, negative binomial seems to be a reasonable choice here
```

####Calculate interaction magnitudes
```{r interaction magnitudes}
intxn <- as.tibble(ft2$fourth)
intxn$resptraits <- rownames(intxn)
intxn2 <- intxn %>% pivot_longer(cols = c(PC1.Woody:Tree_cover)) %>% filter(!value == 0) %>% pivot_wider(id_cols = resptraits)

mean(c(abs(intxn2$Tree_cover),abs(intxn2$Impervious_surf)),na.rm = TRUE)
mean(c(abs(intxn2$PC1.Woody),abs(intxn2$PC2.Herbaceous)), na.rm = TRUE)

mean(c(abs(intxn2$Tree_cover),abs(intxn2$Impervious_surf)),na.rm = TRUE)/mean(c(abs(intxn2$PC1.Woody),abs(intxn2$PC2.Herbaceous)), na.rm = TRUE)
```

###Get top env + trait variables
```{r get top environmental variables}
trait.env.coef1 <- as_tibble(ft2$fourth.corner) %>%  mutate(trait = colnames(Q))

trait.env.coef <- trait.env.coef1 %>% pivot_longer(cols = PC1.Woody:Impervious_surf, names_to = "env.var", values_to = "env.val")

options(scipen=999)
top.95th.val <- quantile(abs(trait.env.coef$env.val), c(.95)) #select the response traits in the top 95th

#select the traits with interaction coefficients in the 95th percentile (most important) to carry forward to the RDA
(trait.env.coef.top <- trait.env.coef %>%
  filter(abs(env.val) >= top.95th.val))

trait.env.coef.top$trait <- str_replace(trait.env.coef.top$trait,"1","")

trait.env.top <- unique(trait.env.coef.top$trait) #these are the traits with the greatest magnitude of trait-environment coefficients
env.top <- unique(trait.env.coef.top$env.var) #these are the traits with the greatest magnitude of trait-environment coefficients

trait.env.top
env.top
```

###Prep data for fourth corner plot
```{r plotting fourth corner results}
fit.4th1 <- as.data.frame(ft2$fourth.corner)
  fit.4th1$Responsetraits <- rownames(fit.4th1)
  
fit.4th1$Responsetraits <- gsub("1", "", fit.4th1$Responsetraits) 
  colnames(Q)
fit.4th <- fit.4th1 %>%
  mutate(Responsetraits = colnames(Q)) %>%
  pivot_longer(-Responsetraits,names_to="Environmentalvariable",values_to = "Envvarvalue")

fit.4th$Responsetraits <- ordered(ordered(fit.4th$Responsetraits, 
                                          levels = c("Structure_nests","Cavity_nests",  "Clutch_size", "Parental_care", "Diet_evenness", "Fruits", "Seeds", "Invertebrates", "Plants", "Body_size", "Bill_length", "Wing_length", "Canopy_foraging", "Ground_foraging", "Midhigh_foraging", "Understory_foraging")))       
fit.4th$Environmentalvariable <- ordered(ordered(fit.4th$Environmentalvariable, 
                                                 levels = c("Impervious_surf", "Tree_cover", "PC1.Woody", "PC2.Herbaceous")))

fit.4th$grtr.zero <- ifelse(round(fit.4th$Envvarvalue, digits = 2) == 0, TRUE, FALSE) 

limit <- max(abs(fit.4th$Envvarvalue)) * c(-1, 1) 

top.95th.val <- quantile(abs(trait.env.coef$env.val), c(.95)) #select the response traits in the top 95th

fit.4th$quant.95 <- ifelse((abs(fit.4th$Envvarvalue) >= top.95th.val), TRUE, FALSE)
```


###Plotting 4th corner interaction coefficients tile plot
```{r 4th corner interaction coefficients tile plot}
dev.off()
dev.new()
(fourth.plot <- ggplot(fit.4th)+
  geom_tile(aes(Responsetraits,Environmentalvariable,fill = Envvarvalue),height = 1, width = 1)+
    geom_tile(data =  fit.4th %>% filter(quant.95 == TRUE), aes(Responsetraits,Environmentalvariable, fill = Envvarvalue), color = "black", size = 0.5, height = 1, width = 1)+
geom_text(data = (fit.4th %>% filter(!grtr.zero == TRUE)),aes(Responsetraits,Environmentalvariable,label = round(Envvarvalue, digits = 3)))+
  scale_fill_gradientn(colors = c("#0072B2","white","#D55E00"), limits = limit)+ ylab("Habitat Variables")+ xlab("Response Traits")+
  theme(axis.text.x = element_text(angle = 45, vjust = 1, hjust=1))+
  coord_flip()+  theme(panel.border = element_rect(colour = "black", fill=NA, size=1))+
  labs(fill="Interaction \nCoefficient")+  theme(legend.position="right")+ theme(panel.background = element_rect(fill = "white", linetype = NULL),panel.grid.major = element_blank(),panel.grid.minor = element_blank()))
```

###*Saving plot PDF
```{r saving figure 1}
#pdf(file = "~/Figure1.pdf",   # The #directory you want to save the file in
    width = 5.5, # The width of the plot in inches
    height = 5.5) # The height of the plot in inches
fourth.plot
dev.off()
```


#Part 3: Response-effect trait correlations
The correlations between response and effect traits create a linkage between habitat and ecosystem services. We perform an abundance-weighted correlation to account for the real representation of traits in the communities we sampled, since not all species (and their response-effect trait combinations) are present at equal abundances in the urban landscape.

##Data Tidying
###prep response traits
```{r prep response traits}
Q.matrix #species x traits matrix

BirdTraits.all.long <- read_excel(all.data, sheet = "BirdTraits.all.long")

birdslist1 <- as.data.frame(Q.matrix$Species)
birdslist <- birdslist1 %>% dplyr::rename(Species = `Q.matrix$Species`)

BirdTraits.all2 <- BirdTraits.all.long %>%
  filter(!TraitType == "C") %>%
  dplyr::select(-TraitType,-TraitGroup) %>%
  pivot_wider(values_from = "Value", names_from = "Trait")

BirdTraits.all2
BirdTraits.all1 <- left_join(birdslist,BirdTraits.all2)
BirdTraits.all1

resp.traits1 <- Q #response traits dataframe
resp.traits1$Species <- (BirdTraits.all1$Species)
resp.traits <- as.tibble(resp.traits1) %>%
  dplyr::select(trait = all_of(trait.env.top)) #%>%

colnames(resp.traits) = (trait.env.top) #reset column names to traits

resp.traits$Species <- (BirdTraits.all1$Species)

resp.traits #gives our final set of response traits to include here.

```

###prep effect traits
```{r prep effect traits}
BirdTraits.Eff <- BirdTraits.all.long %>%
  filter(!TraitType == "R") %>%
  dplyr::select(-TraitType) %>%
  pivot_wider(id_cols= c(Species), values_from = "Value", names_from = "Trait")

BirdTraits.Eff

effect.traits1 <- left_join(birdslist,BirdTraits.Eff, by ="Species")

effect.traits1
```

###prep bird abundances
```{r prep bird abundances}
birdabundances.all <- read_excel(all.data, sheet = "birdabundances.all")

tot.sp.abund <- birdabundances.all1 %>%
  group_by(Species) %>%
  dplyr::summarize(SpAbundance = sum(Abundance50), M.SpAbundance = sum(MeanAbundance50)) 

tot.site.abund <- birdabundances.all1 %>%
  dplyr::select(SiteID,MeanAbundance50,Abundance50) %>%
  group_by(SiteID) %>%
  dplyr::summarize(M.SiteAbundance = sum(MeanAbundance50),
                   SiteAbundance = sum(Abundance50))

tot.site.sp.abund <- birdabundances.all1 %>%
  dplyr::select(SiteID,Species,MeanAbundance50,Abundance50) %>%
  group_by(SiteID,Species) %>%
  dplyr::summarize(M.SiteAbundance = sum(MeanAbundance50),
                   SiteAbundance = sum(Abundance50))
```

##Response-effect trait correlations calculation
```{r response-effect trait correlations} 
library(corrplot)
library(wCorr)

resp.effect.traits.sp.abund <- read_excel(all.data, sheet = "resp.effect.traits.sp.abund")

str(resp.effect.traits.sp.abund)
resp.effect.traits.sp.abund <- ungroup(resp.effect.traits.sp.abund)

resp.effect.traits.sp.abund <- resp.effect.traits.sp.abund %>%
  rename_at(
    vars(ends_with(".x")),
    ~str_replace(., "\\..$","")
  ) %>% 
  select_at(
    vars(-ends_with(".y"))
  )

resp.effect.traits.sp.abund <- as.data.frame(resp.effect.traits.sp.abund)

ResponseTraits1 <- resp.traits %>%
  dplyr::select(-Species)
ResponseTraits <- colnames(ResponseTraits1)

EffectTraits1 <- effect.traits1 %>%
  dplyr::select(-Species)
EffectTraits <- colnames(EffectTraits1)

new.tibble <- NULL
new.tibble <- as.data.frame(expand_grid(ResponseTraits, EffectTraits)) 
new.tibble 

for(i in 1:nrow(new.tibble)){
wcors = weightedCorr(
                    y = resp.effect.traits.sp.abund[,new.tibble[i,"EffectTraits"]],
                    x = resp.effect.traits.sp.abund[,new.tibble[i,"ResponseTraits"]], 
                    method = "pearson", 
                    weights = resp.effect.traits.sp.abund$M.SpAbundance #weight by mean abundance for each species at each site across all sampling occasions. Same weighting as RDA and 4th corner
                    )
cors = cor(
                    y = resp.effect.traits.sp.abund[,new.tibble[i,"EffectTraits"]],
                    x = resp.effect.traits.sp.abund[,new.tibble[i,"ResponseTraits"]], 
                    method = "pearson"
                    )

new.tibble[i,"wcors"] <- wcors
new.tibble[i,"cors"] <- cors
}

new.tibble["wcors"] <- round(new.tibble[,"wcors"], digits=10) 

head(new.tibble) #check output
```

###testing R-E Correlatations for significance
```{r significance testing of Pearson's cors}
#bonferroni-corrected alpha level:
a = 0.05 #initial alpha level
tests = nrow(new.tibble) #number of tests
bc.a <- a/tests

criticalt <- qt(p=bc.a, df=46, lower.tail=FALSE)

new.tibble.sig1 <- new.tibble %>%
  mutate(twcors = abs(wcors/(sqrt(1-(wcors^2)))) * (sqrt(46-2)), wcor.sig = twcors >= criticalt)

```

###Plotting R-E trait correlation matrix
```{r plotting correlation tile plots}
new.tibble.sig1$EffectTraits <- ordered(new.tibble.sig1$EffectTraits, c("Black_plumage", "Cool_plumage", "Dull_plumage", "Warm_plumage", "Crest_presence", "Frequency_range", "Syllable_count", "Diet_evenness", "Fruits", "Invertebrates", "Seeds","Tail_length","Bill_length","Body_size"))

new.tibble.sig1$Trait <- new.tibble.sig1$EffectTraits

new.tibble.sig <- left_join(as_tibble(new.tibble.sig1), BirdTraits.all.long %>% dplyr::select(TraitGroup, Trait) %>% distinct())

dev.off()
dev.new()
(retraits.plot <- ggplot(new.tibble.sig) +
  geom_tile(aes(y = EffectTraits, x = ResponseTraits, fill = wcors),height = 1, width = 1)+
  scale_fill_gradient2(low = "#00798c", mid = "white", high = "#edae49")+
    geom_tile(data = (new.tibble.sig %>% filter(wcor.sig == TRUE)), aes(y = EffectTraits, x = ResponseTraits, fill = wcors), color = "black",height = 1, width = 1, size = 0.5)+
geom_text(aes(y = EffectTraits, x = ResponseTraits, label=round(wcors, digits = 3)))+ 
    theme(axis.text.x = element_text(angle = 45, vjust = 1, hjust=1)) + theme(panel.border = element_rect(colour = "black", fill=NA, size=1))+ 
theme(panel.background = element_rect(fill = "white", linetype = NULL),panel.grid.major = element_blank(),panel.grid.minor = element_blank())+
  labs(fill = "Weighted \nCorrelation",y = "Effect Traits", x = "Response Traits") + facet_wrap(~TraitGroup, scales = "free_y", ncol = 2))
```

###*Saving plot PDF
```{r saving figure 2}
#pdf(file = "~/Figure2.pdf",   # The #directory you want to save the file in
    width = 8, # The width of the plot in inches
    height = 5) # The height of the plot in inches
retraits.plot
dev.off()
```


#Part 4: RDA for community-summed effect traits
In this analysis, we employ a redundancy analysis to explain how community-summed effect trait values at each site vary in relation to the local and landscape habitat present. This indicates associations between habitat and summed effect traits.

```{r loading data pt3}
library(tidyverse)

sumtraits2 <- read_excel(all.data, sheet = "cset.sites.traits.all")

veg.site.scores <- read_excel(all.data, sheet = "veg.site.scores")

pcveg.data2 <- read_excel(all.data, sheet = "PCVegData")

sumtraits1 <- sumtraits2 %>%  filter(!SiteID == "L-14")
env <- left_join(pcveg.data2,veg.site.scores)
```

##Data Wrangling
###Community-summed effect trait (CSET) data
```{r data wrangling pt3}
(sumtraits <- sumtraits1 %>%
    transmute(SiteID = SiteID,
           Diet_evenness = CSET.Evenness,
           Invertebrates = CSET.Diet_Inv,
           Seeds = CSET.Diet_Seed,
           Fruits = CSET.Diet_Fruit,
           Frequency_range = CSET.DeltaFreqHz,
           Syllable_count = CSET.SylCount,
           Body_size = CSET.BodyMass_Value,
           Warm_plumage = CSET.warm,
           Tail_length = CSET.TailBodyRatio,
           Bill_length = CSET.BillBodyRatio,
           Cool_plumage = CSET.cool,
           Black_plumage = CSET.black,
           Dull_plumage = CSET.dull,
           Crest_presence = CSET.Present)) 
head(sumtraits)
```


###Centering and scaling traits
```{r centering and scaling community-summed trait values}
#centering and scaling
library(caret) #for centering and scaling the data

#traits
preObj <- preProcess(sumtraits, method=c("center", "scale"))
traits <- predict(preObj, sumtraits[,])

summary(traits) #seem to all be looking pretty good now. 

env7 <- env %>%  mutate(PC1.Woody = PC1,
PC2.Herbaceous = PC2) %>% dplyr::select(-PC1,-PC2,-PC3,-PC4,-PC5,-PC6,-DetailType)

#env vars
preObj <- preProcess(env7, method=c("center", "scale"))
env.rda <- predict(preObj, env7[,])

traits.env <- cbind(env.rda[,-1], traits[,-c(1)])
```


##Running RDA
```{r running RDA}
library(vegan) #load the vegan package, which is used for the RDA

spe.rda <- 
  rda(traits[,-c(1)] ~ Impervious_surf + Tree_cover + PC1.Woody + PC2.Herbaceous,  data = env.rda)

summary(spe.rda)
```
###RDA stats
```{r getting RDA stats}
rda4 <- rda(traits[,-c(1)]) 
envfit(rda4,env = env.rda[,c("Tree_cover","Impervious_surf","PC1.Woody","PC2.Herbaceous")])

#Checking for multicollinearity
vif.cca(spe.rda)

# Canonical coefficients from the rda object
coef(spe.rda)

anova(spe.rda,permutations= 1000)

# Unadjusted R^2 retrieved from the rda object
(R2 <- RsquareAdj(spe.rda)$r.squared)
(R2adj <- RsquareAdj(spe.rda)$adj.r.squared)

#Permutation test to identify significant axis (axes)
anova.cca(spe.rda,permutations= 1000, by = "axis")

var.expl <- spe.rda$CCA$eig/spe.rda$tot.chi #create an object that contains the % variation explained for each of the RDA axes
var.expl

screeplot(spe.rda)

#Are any of the habitat variables significant?
anova.cca(spe.rda,permutations= 1000, by = "terms")

```

##RDA Biplot
###Prepping data
```{r prepping data for trait family biplots}
library(vegan)
scor = scores(spe.rda, display=c("sites", "species", "bp"), scaling=3)

species_centroids <- data.frame(scor$species)
species_centroids
species_centroids$trait_name <- rownames(species_centroids) 

sites_centroids <- data.frame(scor$sites)
sites_centroids
sites_centroids$SiteID <- env.rda$SiteID 

#env var arrows
env_arrows <- data.frame(scor$biplot)
env_arrows
env_arrows$env_var <- rownames(env_arrows) 
env_arrows

#scaling for arrows
basplot <- plot(spe.rda)
mult <- attributes(basplot$biplot)$arrow.mul 

```

###Subsetting data for trait family biplots
```{r subsetting by trait family and trait type}
species_centroids$traitfam <- c("Diet","Diet","Diet","Diet",
                         "Acoustic","Acoustic",
                         "Visual","Visual", "Visual","Visual","Visual","Visual", "Visual","Visual"
                         )
species_centroids$traittype <- c("Response+Effect","Response+Effect","Response+Effect","Effect","Effect","Effect",
                          "Response+Effect","Effect","Effect","Effect","Effect","Effect", "Effect","Effect")
species_centroids$service <- c("Diet Generality", "Invertivory", "Diet_granivory", "Frugiveroy", "Acoustic", "Acoustic", "Visual_shape", "Visual_plumage", "Visual_shape", "Visual_shape", "Visual_plumage", "Visual_plumage", "Visual_plumage", "Visual_shape")
acoustic <- subset(species_centroids,traitfam=='Acoustic')
visual <- subset(species_centroids,traitfam=='Visual')
diet.forage <- subset(species_centroids,traitfam=='Diet')
resp.only <- subset(species_centroids,traittype=='Response' | traittype== 'Response+Effect')
```

###RDA Biplot (all variables)
```{r ggplot of all families}
library(ggrepel)
species_centroids$traitfam <- as.factor(species_centroids$traitfam)
species_centroids$traitfam <- ordered(species_centroids$traitfam,c("Diet", "Acoustic", "Visual"))
dev.off()
dev.new()
(RDA_plot <- ggplot(env_arrows, aes(x = RDA1, y = -RDA2))+
    #Add arrows for environmental variables 
    geom_segment(data = (env_arrows %>% dplyr::filter(env_var == "log.AreaM2"|env_var ==  "Tree_cover" | env_var ==  "Impervious_surf" | env_var == "PC1.Woody" | env_var == "PC2.Herbaceous")), aes(x = 0, xend = mult * RDA1,y = 0, yend = mult * -RDA2), arrow = arrow(length = unit(0.25, "cm")), colour = "gray40") + 
    geom_text_repel(data = (env_arrows %>% dplyr::filter(env_var == "log.AreaM2"|env_var ==  "Tree_cover" | env_var ==  "Impervious_surf" | env_var == "PC1.Woody" | env_var == "PC2.Herbaceous")),aes(x= (mult + mult/10) * RDA1, y = (mult + mult/10) * -RDA2, label = env_var), size = 3.5,color = "black")+ 
    #Add points for Effect Traits
    geom_point(data = species_centroids) + geom_text_repel(data = species_centroids, aes(label = trait_name), vjust = -0.5,size = 3)+
    
    #Add miscellaneous plot parameters
    geom_vline(aes(xintercept = 0), linetype = "dashed",color = "black",alpha = 0.5) +
    geom_hline(aes(yintercept = 0), linetype = "dashed",color = "black",alpha = 0.5) +
    labs(color = "Ecosystem Service")+
    scale_color_manual(values=c("#E69F00", "#56B4E9", "#009E73", "#F0E442", "#0072B2", "#D55E00", "#CC79A7"))+
    xlab(paste0("RDA1  ", round(var.expl[1]*100,2), "%"))+ ylab(paste0("RDA2  ", round(var.expl[2]*100,2), "%"))+
    
    #Set theme elements
    theme(legend.position = "top") + theme(panel.border =  element_rect(colour = "black", fill=NA, size=0.5))+
    theme(panel.background = element_rect(fill = "gray93", linetype = NULL),panel.grid.major = element_blank(),panel.grid.minor = element_blank())
)

```

###*Saving plot PDF
```{r saving figure 2}
#pdf(file = "~/Figure3.pdf",   # The #directory you want to save the file in
    width = 6, # The width of the plot in inches
    height = 6) # The height of the plot in inches
RDA_plot
dev.off()
```

###Plot trait loadings
```{r plotting trait loadings}
species_centroids

library(ggpubr)
library(cowplot)

(trait.rda1 <- ggplot(species_centroids %>% filter(!traitfam == "NA"))+
  geom_bar(aes(x = reorder(trait_name,RDA1),y = RDA1,fill = traitfam),stat = "identity")+
  coord_flip()+
  xlab("Trait")+
  guides(fill = FALSE)+
  ylab(paste0("RDA1  ", round(var.expl[1]*100,2), "%"))+
     theme_pubclean()+
 theme(panel.border =  element_rect(colour = "black", fill=NA, size=0.5))+
  theme(panel.background = element_rect(fill = "gray93", linetype = NULL),panel.grid.major = element_blank(),panel.grid.minor = element_blank())+

  scale_fill_manual(values = c("#0072B2","#009E73","#D55E00")))

(trait.rda2 <- ggplot(species_centroids %>% filter(!traitfam == "NA"))+
    geom_bar(aes(x = reorder(trait_name,-RDA2),y = -RDA2, fill = traitfam),stat = "identity")+
    coord_flip()+
    xlab("")+
    theme_pubclean()+
    labs(fill = "Trait Family")+
    ylab(paste0("RDA2  ", round(var.expl[2]*100,2), "%"))+
    theme(legend.position = "right")+
    theme(panel.border =  element_rect(colour = "black", fill=NA, size=0.5))+
    theme(panel.background = element_rect(fill = "gray93", linetype = NULL),panel.grid.major = element_blank(),panel.grid.minor = element_blank())+
    scale_fill_manual(values = c("#0072B2","#009E73","#D55E00")))

plot_grid(trait.rda1, trait.rda2, labels = c('a)', 'b)'), label_size = 12)
```


#Part 5a: Ecosystem Service Calculations 
Here we model ecosystem service supply based on expected relationships between the effect traits underlying the services.

```{r modeling site level ESs} 
preObj <- preProcess(sumtraits, method=c("scale"))
traits.scaled <- predict(preObj, sumtraits[,])

traits.es.scaled <- traits.scaled %>%
  pivot_longer(Diet_evenness:Crest_presence, values_to =  "value", names_to = "variable") %>%
  mutate(value = value+abs(min(value))) %>% 
  pivot_wider(values_from = "value", names_from = "variable")

head(traits.es.scaled) 

env
summary(env)

traits.env.es.scaled <- left_join(env, traits.es.scaled)

head(traits.env.es.scaled)
```

##Calcuating Summed Ecosystem Services
```{r calculate summed service levels}

(sumtraits.env.ES <- traits.env.es.scaled %>%
   mutate(Acoustic_aesthetics = Syllable_count + Frequency_range,
          Plumage_aesthetics = (Warm_plumage + Cool_plumage) - (Black_plumage + Dull_plumage),
          Shape_aesthetics = (Crest_presence + Tail_length) - (Body_size + Bill_length),
          Invertivory = (Invertebrates),
          Diet_evenness = (Diet_evenness),
          Frugivory = (Fruits),
          Granivory = (Seeds),
          PC1.Woody = PC1,
          PC2.Herbaceous = PC2
   ) %>%
   dplyr::select(SiteID,
                 DetailType,
                 Impervious_surf,
                 Tree_cover,
                 PC1.Woody, PC2.Herbaceous,
                 Acoustic_aesthetics,
                 Plumage_aesthetics,
                 #Visual_plumage_Div,
                 Shape_aesthetics,
                 Diet_evenness,
                 Invertivory,
                 Frugivory,
                 Granivory
   ))

sumtraits.env.ES.long <- sumtraits.env.ES %>%
  pivot_longer(Acoustic_aesthetics:Granivory,names_to = "EcosystemService", values_to = "TraitVal")

head(sumtraits.env.ES)

```


#Part 5b: Predicting ecosystem service supply along habitat gradients
Here we used linear regressions to model and test the significance of relationships between local and landscape habitat gradients and the modeled ecosystem service values we calculated.

###Significance Tests
```{r significance tests for ES-Impervious relationships}
sites_centroids

sumtraits.env.ES1 <- as.data.frame(sumtraits.env.ES)
varlist <- c("Acoustic_aesthetics", "Plumage_aesthetics", "Shape_aesthetics", "Diet_evenness", "Invertivory", "Frugivory", "Granivory")

sumtraits.sites.rda <- left_join(sites_centroids,sumtraits.env.ES1)

models <- lapply(varlist, function(x) {glm(substitute(i ~ Impervious_surf, list(i = as.name(x))), data = sumtraits.sites.rda)})
models.sum <- lapply(models, summary)

names(models.sum) <- varlist
```

###Get mod stats
```{r get mod stats}
getpval <- function(models.sum) {
  p.val1 <- models.sum$coefficient[8]
  R2 <- (1-(models.sum[["deviance"]])/(models.sum[["null.deviance"]]))
  c(p.val= p.val1,R2 = R2)
}

getmodstats <- function(models.sum) {
  p.val1 <- models.sum$coefficient[8]
  beta <- models.sum$coefficient[2]
  SE <- models.sum$coefficient[4]
  R2 <- (1-(models.sum[["deviance"]])/(models.sum[["null.deviance"]]))
  c(p.val= p.val1,R2 = R2, beta = beta, SE = SE)
  }

model.outputs2 <- as.data.frame(sapply(models.sum, getmodstats))

model.outputs2$stat <- rownames(model.outputs2)
model.outputs2$envvar <- "ImperviousSurface"

model.outputs1 <- model.outputs2 %>%
  pivot_longer(cols = c(1,2,3,4,5,6,7), names_to = "var") %>%
  pivot_wider(values_from = value, names_from = stat) %>%
  mutate(p.val1 = p.val) %>%
  mutate(p.val3 = round(p.val1, digits = 3),
         SE = SE,
         SD = sqrt(60)*SE,
         SE = round(as.numeric(SE),digits = 4),
         CI.95.pos = beta+1.960*(SD/sqrt(60)),
         CI.95.neg = beta-1.960*(SD/sqrt(60)),
         beta = round(as.numeric(beta),digits = 4)
         )

model.outputs1$p.val <- ifelse(model.outputs1$p.val3 == "0", "<0.0001", round(model.outputs1$p.val3, digits = 3))

model.outputs1$p.sig <- ifelse(model.outputs1$p.val1 <= (0.05/nrow(model.outputs1)), TRUE, FALSE) #bonferroni corrected critical value

model.outputs <- model.outputs1 %>%
  dplyr::select(-p.val1, -p.val3)
```


##Plotting betas
```{r plotting betas for summed Ecosystem Services}
model.outputs

model.outputs$EcosystemService <- ordered(model.outputs$var, levels = c("Frugivory","Diet_evenness","Granivory","Invertivory","Shape_aesthetics","Plumage_aesthetics","Acoustic_aesthetics"))

(betas.plot <- ggplot(model.outputs)+
  geom_point(aes(y = beta, x = EcosystemService, shape = p.sig))+
  scale_shape_manual(values=c(1,16))+
  geom_errorbar(aes(y = beta, x = EcosystemService, ymin = CI.95.neg, ymax = CI.95.pos),width = 0.0)+
  geom_hline(yintercept = 0,linetype='dotted')+
  coord_flip()+
    theme(legend.position = "none")+
  theme(panel.border =  element_rect(colour = "black", fill=NA, size=0.5))+
  theme(panel.background = element_rect(fill = "gray93", linetype = NULL),panel.grid.major = element_blank(),panel.grid.minor = element_blank())+
  labs(y = "Impervious surface (betas, 95% CI)", x = "Ecosystem Service"))
```

####*Saving plot PDF
```{r saving figure 2}
#pdf(file = "~/Figure4.pdf",   # The #directory you want to save the file in
    width = 5, # The width of the plot in inches
    height = 4) # The height of the plot in inches
betas.plot
dev.off()
```
````
