## Supplementary Tables and Figures for "A functional trait approach reveals the effects of landscape context on ecosystem services provided by urban birds"

**Supporting Information**

T. M. Swartz, J.M. Gleditsch, and J. E. Behm. 2022. Bird functional traits and ecosystem services

Supporting Tables and Figures specific to the analysis of bird functional traits and ecosystem services in urban green spaces sampled in summer 2019 in Philadelphia County, Pennsylvania, USA.

**Supporting Tables**

**Table S1.** GPS Coordinates in decimal degrees of Philadelphia urban green spaces where bird communities and habitat were sampled in 2019 (datum WGS 1984).

| SiteID | Green Space Type | Latitude | Longitude |
|---|---|---|---|
| L-01 | Ungreened vacant lot | 40.04695 | -75.1563 |
| L-02 | Ungreened vacant lot | 39.98949 | -75.1651 |
| L-03 | Ungreened vacant lot | 40.02252 | -75.0668 |
| L-06 | Ungreened vacant lot | 39.973 | -75.1606 |
| L-07 | Ungreened vacant lot | 40.01262 | -75.0828 |
| L-09 | Ungreened vacant lot | 39.96833 | -75.2271 |
| L-11 | Ungreened vacant lot | 40.03341 | -75.1599 |
| L-12 | Ungreened vacant lot | 39.909906 | -75.245012 |
| L-16 | Ungreened vacant lot | 40.03301 | -75.1652 |
| L-17 | Ungreened vacant lot | 40.03149 | -75.227 |
| P-05 | Recreation park | 39.99064 | -75.0972 |
| P-06 | Recreation park | 39.92122 | -75.1504 |
| P-07 | Recreation park | 39.98603 | -75.1516 |
| P-08 | Recreation park | 39.98427 | -75.1116 |
| P-11 | Recreation park | 39.99425 | -75.1235 |
| P-12 | Recreation park | 39.97772 | -75.1787 |
| P-13 | Recreation park | 39.98109 | -75.1929 |
| P-18 | Recreation park | 40.042456 | -74.992505 |
| P-31 | Recreation park | 40.01234 | -75.0969 |
| P-32 | Recreation park | 40.05162 | -75.0026 |
| P-35 | Recreation park | 40.04898 | -75.1239 |
| P-41 | Recreation park | 40.06786 | -74.9912 |
| P-15 | Forest preserve | 40.05903 | -75.2113 |
| P-16 | Forest preserve | 40.080283 | -75.231844 |
| P-20 | Forest preserve | 40.06924 | -75.2311 |
| P-21 | Forest preserve | 39.97709 | -75.2589 |
| P-24 | Forest preserve | 40.00142 | -75.1938 |
| P-25 | Forest preserve | 40.06029 | -75.0369 |
| P-26 | Forest preserve | 39.982099 | -75.257429 |
| P-27 | Forest preserve | 40.03401 | -75.2306 |
| P-28 | Forest preserve | 40.051995 | -75.213221 |
| P-33 | Forest preserve | 39.989135 | -75.194037 |

| | | | |
|---|---|---|---|
| P-40 | Forest preserve | 40.022545 | -75.201045 |
| P-01 | Plaza | 39.9538 | -75.2216 |
| P-02 | Plaza | 39.92049 | -75.1832 |
| P-03 | Plaza | 39.93925 | -75.1556 |
| P-04 | Plaza | 39.92767 | -75.1516 |
| P-09 | Plaza | 40.00908 | -75.1383 |
| P-10 | Plaza | 39.94699 | -75.1527 |
| P-22 | Plaza | 40.05164 | -74.9805 |
| P-23 | Plaza | 40.06715 | -75.2031 |
| P-29 | Plaza | 39.96351 | -75.1782 |
| P-30 | Plaza | 39.943728 | -75.142754 |
| P-34 | Plaza | 39.9473 | -75.2097 |
| P-36 | Plaza | 40.01809 | -75.1877 |
| L-19 | Greened vacant lot | 40.02413 | -75.1647 |
| L-20 | Greened vacant lot | 39.96564 | -75.2004 |
| L-21 | Greened vacant lot | 39.98583 | -75.17 |
| L-22 | Greened vacant lot | 40.05365 | -75.1828 |
| L-23 | Greened vacant lot | 40.00029 | -75.154 |
| L-24 | Greened vacant lot | 39.974621 | -75.224479 |
| L-26 | Greened vacant lot | 39.98527 | -75.1465 |
| L-27 | Greened vacant lot | 40.00651 | -75.1577 |
| L-28 | Greened vacant lot | 40.05131 | -75.1815 |
| L-29 | Greened vacant lot | 39.98094 | -75.1795 |
| L-30 | Greened vacant lot | 39.96912 | -75.2234 |
| L-31 | Greened vacant lot | 40.04393 | -75.158 |
| L-32 | Greened vacant lot | 39.98285 | -75.1434 |
| L-33 | Greened vacant lot | 40.03729 | -75.1655 |
| L-34 | Greened vacant lot | 40.03579 | -75.1626 |

**Table S2.** The five types of green spaces where our 60 study sites were located

| **Green Space Type** | **Management** | **Habitat Structure** | **Count (N)** |
|---|---|---|---|
| Ungreened vacant Lot | Many are neglected, mowing and vegetation management may be infrequent/absent | Consists of spontaneous, naturally-colonizing vegetation | 10 |
| Greened vacant lot | Mowing and vegetation occur regularly through Philadelphia's vacant lot program (see South er al. 2018) | Some trees along edges, grassy lawns | 15 |
| Plaza | Regular mowing and vegetation maintenance | Usually with large shade trees, open lawns, and limited understory vegetation | 12 |
| Recreation park | Regular mowing and vegetation maintenance | Some trees along edges, well-maintained grassy lawns recreation activities | 12 |
| Forest preserve | Largely unmanaged, but some tree planting and invasive species management may occur | Natural vegetation typical of deciduous forest, sometimes with dense understory | 11 |

**Table S3.** List of bird recording files downloaded from www.xeno-canto.org and used in the song analysis procedure. XCID is the official www.xeno-canto.org record number.

| Species | EnglishName | Recordist | XCID | RecordingID | Type | Mimic |
|---|---|---|---|---|---|---|
| AMGO | American Goldfinch | Dan Lane | XC179086 | AMGO_XC179086 - American Goldfinch - Spinus tristis | Song | Y |
| BGGN | Blue-gray Gnatcatcher | Paul Driver | XC138062 | BGGN_XC138062 - Blue-grey Gnatcatcher - Polioptila caerulea | Call | Y |
| BLJA | Blue Jay | Orlando Jarquan G. | XC385026 | BLJA_XC385026 - Blue Jay - Cyanocitta cristata | Call | Y |
| EUST | European Starling | Daniel Parker | XC162972 | EUST_XC162972 - Common Starling - Sturnus vulgaris | Song | Y |
| GRCA | Gray Catbird | Paul Driver | XC248135 | GRCA_XC248135 - Grey Catbird - Dumetella carolinensis | Song | Y |
| HOFI | House Finch | Paul Driver | XC169187 | HOFI_XC169187 - House Finch - Haemorhous mexicanus | Song | Y |
| NOMO | Northern Mockingbird | Larry Simms | XC468619 | NOMO_XC468619 - Northern Mockingbird - Mimus polyglottos | Song | Y |
| AMCR | American Crow | Paul Driver | XC196757 | AMCR_XC196757 - American Crow - Corvus brachyrhynchos | Call | N |
| AMRE | American Redstart | Jerald R | XC245807 | AMRE_XC245807 - American Redstart - Setophaga ruticilla | Song | N |
| AMRO | American Robin | Rolf A. de By | XC469605 | AMRO_XC469605 - American Robin - Turdus migratorius | Song | N |
| BAOR | Baltimore Oriole | Paul Marvin | XC217799 | BAOR_XC217799 - Baltimore Oriole - Icterus galbula | Song | N |
| BCCH | Black-capped Chickadee | Jim Berry | XC469109 | BCCH_XC469109 - Black-capped Chickadee - Poecile atricapillus | Song | N |
| BHCO | Brown-headed Cowbird | Matt Wistrand | XC312718 | BHCO_XC312718 - Brown-headed Cowbird - Molothrus ater | Song | N |
| BLPW | Blackpoll Warbler | Dan Lane | XC180072 | BLPW_XC180072 - Blackpoll Warbler - Setophaga striata | Song | N |
| CACH | Carolina Chickadee | Dan Lane | XC179084 | CACH_XC179084 - Carolina Chickadee - Poecile carolinensis | Song | N |
| CARW | Carolina Wren | Amy Davis | XC141908 | CAWR_XC141908 - Carolina Wren - Thryothorus ludovicianus | Song | N |
| CEDW | Cedar Waxwing | William Whitehead | XC476658 | CEDW_XC476658 - Cedar Waxwing - Bombycilla cedrorum | Song | N |
| CHSP | Chipping Sparrow | Daniel Lane | XC100725 | CHSP_XC100725 - Chipping Sparrow - Spizella passerina | Call | N |

| | | | | | | |
|---|---|---|---|---|---|---|
| COGR | Common Grackle | Richard E. Webster | XC184987 | COGR_XC184987 - Common Grackle - Quiscalus quiscula versicolor | Song | N |
| COHA | Cooper's Hawk | Paul Driver | XC70471 | COHA_XC70471 - Cooper's Hawk - Accipiter cooperii | Call | N |
| COYE | Common Yellowthroat | Daniel Lane | XC101432 | COYE_XC101432 - Common Yellowthroat - Geothlypis trichas | Song | N |
| DOWO | Downy Woodpecker | Paul Driver | XC248137 | DOWO_XC248137 - Downy Woodpecker - Dryobates pubescens | Call | N |
| EAKI | Eastern Kingbird | Paul Driver | XC140649 | EAKI_XC140649 - Eastern Kingbird - Tyrannus tyrannus | Song | N |
| EAWP | Eastern Wood-pewee | Dan Lane | XC244482 | EAWP_XC244482 - Eastern Wood Pewee - Contopus virens | Song | N |
| FICR | Fish Crow | David Eberly | XC252720 | FICR_XC252720 - Fish Crow - Corvus ossifragus | Call | N |
| GCFL | Great Crested Flycatcher | | XC139093 | GCFL_XC139093 - Great Crested Flycatcher - Myiarchus crinitus | Song | N |
| HOSP | House Sparrow | Dan Lane | XC180085 | HOSP_XC180085 - House Sparrow - Passer domesticus | Call | N |
| HOWR | House Wren | Dan Lane | XC179088 | HOWR_XC179088 - House Wren - Troglodytes aedon | Song | N |
| NOCA | Northern Cardinal | Daniel Lane | XC101437 | NOCA_XC101437 - Northern Cardinal - Cardinalis cardinalis | Song | N |
| NOFL | Northern Flicker | Paul Driver | XC170551 | NOFL_XC170551 - Northern Flicker - Colaptes auratus | Call | N |
| RBWO | Red-bellied Woodpecker | Daniel Lane | XC101438 | RBWO_XC101438 - Red-bellied Woodpecker - Melanerpes carolinus | Call | N |
| REVI | Red-eyed Vireo | Patrick Blake | XC324883 | REVI_XC324883 - Red-eyed Vireo - Vireo olivaceus | Song | N |
| ROPI | Rock Pigeon | Paul Marvin | XC460854 | ROPI_XC460854 - Rock Dove - Columba livia | Call | N |
| RTHU | Ruby-throated Hummingbird | Russ Wigh | XC505916 | RTHU_XC505916 - Ruby-throated Hummingbird - Archilochus colubris | Call | N |
| RWBL | Red-winged Blackbird | Paul Driver | XC141763 | RWBL_XC141763 - Red-winged Blackbird - Agelaius phoeniceus | Song | N |
| SCTA | Scarlet Tanager | David Eberly | XC179560 | SCTA_XC179560 - Scarlet Tanager - Piranga olivacea | Song | N |
| SOSP | Song Sparrow | Paul Driver | XC189286 | SOSP_XC189286 - Song Sparrow - Melospiza melodia | Song | N |
| SWTH | Swainson's Thrush | Matthew R. Halley | XC489007 | SWTH_XC489007 - Swainson's Thrush - Catharus ustulatus swainsoni | Song | N |
| TUTI | Tufted Titmouse | Rolf A. de By | XC469582 | TUTI_XC469582 - Tufted Titmouse - Baeolophus bicolor | Song | N |
| VEER | Veery | Don Jones | XC1228 | VEER_XC1228 - Veery - Catharus fuscescens | Song | N |

| | | | | | | |
|---|---|---|---|---|---|---|
| WAVI | Warbling Vireo | William Whitehead | XC480679 | WAVI_XC480679 - Warbling Vireo - Vireo gilvus | Song | N |
| WBNU | White-breasted Nuthatch | Paul Driver | XC170552 | WBNU_XC170552 - White-breasted Nuthatch - Sitta carolinensis | Song | N |
| WIFL | Willow Flycatcher | Dan Lane | XC134749 | WIFL_XC134749 - Willow Flycatcher - Empidonax traillii | Song | N |
| WOTH | Wood Thrush | Paul Driver | XC139092 | WOTH_XC139092 - Wood Thrush - Hylocichla mustelina | Song | N |

**Table S4.** Equations used to calculate the ecosystem service score for seven services based on community-summed effect trait value (CSET). The sources listed provide details on the directionality of trait-service relationships represented in the models.

| Ecosystem Service | Model | Explanation of ecosystem service relationship | Sources |
|---|---|---|---|
| Plumage_aesthetics | $[CSET_{warm} + CSET_{cool}] - [CSET_{dull} + CSET_{black}]$ | • Warm (red, orange, yellow) and cool (blue, purple, green) plumage hues are more aesthetically appealing than<br>• Black and dull (gray, brown, tan) plumage hues are less aesthetically appealing than other hues | (Frynta et al. 2010, Lišková and Frynta 2013, Echeverri et al. 2019a, Zoeller et al. 2020) |
| Shape_aesthetics | $[CSET_{crest} + CSET_{tail}] - [CSET_{bill} + CCSET_{size}]$ | • Smaller birds are more appealing than large ones.<br>• Birds with longer bills relative to body size are less appealing<br>• Birds with longer tails relative to body size are more appealing<br>• Crests are an appealing feature of birds | (Bjerke and Østdahl 2004)<br>(Frynta et al. 2010)<br>(Lišková and Frynta 2013, Echeverri et al. 2019a)<br>(Echeverri et al. 2019a) |
| Acoustic_aesthetics | $CSET_{syllable} + CSET_{delta}$ | • Songs containing more syllables or with greater changes in pitch could be perceived as more interesting and acoustically appealing | (Blackburn et al. 2014) |
| Invertivory | $[CSET_{invertivory}]$ | • Invertivores can contribute to control of pest arthropods | (Whelan et al. 2015) |
| Frugivory | $[CSET_{frugivory}]$ | • Frugivores can facilitate seed dispersal | (Whelan et al. 2015) |
| Granivory | $[CSET_{seed}]$ | • Granivores may control weeds and promote plant diversity by depredating seeds, may also consume carbohydrate-rich littered food refuse | (Whelan et al. 2015) |
| Diet_generality | $[CSET_{evenness}]$ | • Diet generalists may provide a range of trophic-based services, as well as consume a variety of littered food refuse | |

**Supporting Figures**

**Figure S1.** Principal components (PC) ordination biplot of 60 sites in Philadelphia based on local-scale habitat composition and structure. Sites are represented as points grouped by green space type (colors). Convex hulls are drawn around all sites of the same type. Habitat variables constituting the principal components are represented as labelled vector arrows. Percent of total variance explained by each principal component is displayed on each axis.

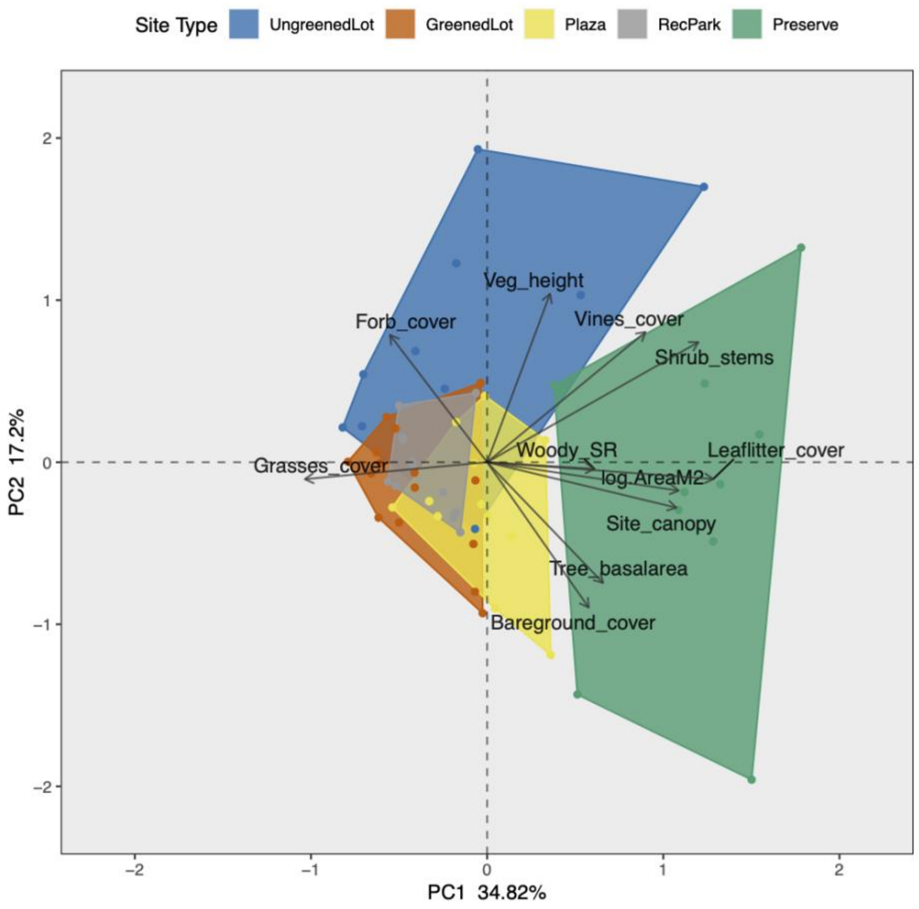

**Figure S2.** Loadings for habitat variables along PC1 and PC2.

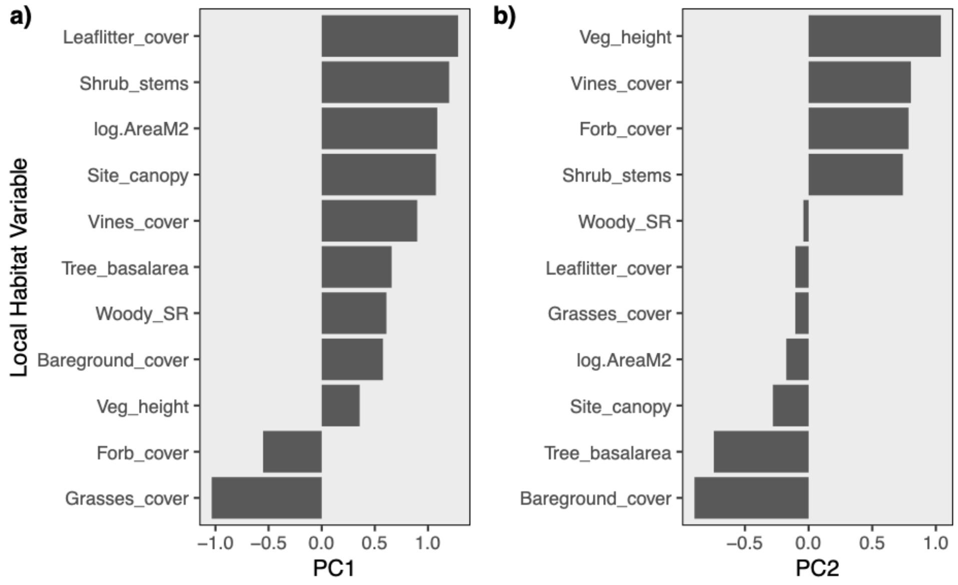
